# Temporal proteomic and PTMomic atlas of cerebral organoid development

**DOI:** 10.1101/2024.09.03.610941

**Authors:** Sofie B. Elmkvist, Helle Bogetofte, Pia Jensen, Lene A. Jakobsen, Jesper F. Havelund, Matias Ryding, Jonathan Brewer, Nils J. Færgeman, Madeline A. Lancaster, Martin R. Larsen

## Abstract

Cerebral organoids (CBOs) are generated from pluripotent stem cells that undergo neuroectoderm specification and neuronal differentiation in three dimensions. The developing neurons in CBOs migrate and self-organize into cerebral cortex-like layers, mimicking human brain development. CBOs develop according to intrinsic signaling mechanisms and offer unique insights into mechanisms of early human brain development. This process requires coordinated spatiotemporal regulation of protein expression and function, where the latter can be achieved by post-translational modifications (PTMs) on proteins. Despite the importance of proteins in brain development and function, profiling of protein abundance and the involvement of PTMs in CBO development remain underexplored. To gain insight into protein and PTM abundance in CBOs, we performed a high-resolution temporal analysis of CBOs up to day 200 using proteomics, PTMomics and metabolomics. We quantified more than 9,300 proteins and various neurodevelopmentally relevant PTMs (including phosphorylation, lysine acetylation, sialylated N-glycosylation, and cysteine modifications). We demonstrate that protein abundance and dynamic PTMs show significant temporal changes during CBO development related to neuronal differentiation and energy metabolism, whereas calcium signaling is mainly regulated by dynamic PTMs. We further show that synaptic protein content correlated with neurotransmitter levels, and we detected astroglia beyond day 100. Lastly, comparative analysis showed proteomic similarities between CBOs and human fetal brain tissue, supporting the physiological relevance of CBOs. Overall, our study presents a temporal atlas of protein and PTM abundance in CBOs and provides a valuable resource for studying neurodevelopment in neural organoids.

## Introduction

Cerebral organoids (CBOs), also known as unguided neural organoids, are three-dimensional structures derived from pluripotent stem cells (PSCs) that mimic early brain development [1-3]. PSCs cultured in suspension undergo neuroectoderm specification and neuronal differentiation. During the process, newly born neurons self-organize into layers resembling the architecture of the developing cerebral cortex. CBOs are cultured without the use of guidance molecules and develop according to intrinsic temporal cellular signaling programs and gene regulatory networks, thus offering unique insights into human brain development [1, 4, 5]. Gene regulatory networks are essential for cell fate determination and differentiation during brain development. These networks rely on the coordinated expression and regulation of proteins through transcription factors, epigenetic modifiers, splicing, and post-translational modifications (PTMs) [6-11]. Proteins play key roles in most cellular functions and signaling pathways, including neuronal migration, axon guidance, and neurotransmission, which are critical processes for brain development and function [12, 13]. The regulation of protein expression is closely linked with changes in gene activity, and the regulation of PTMs on proteins enables dynamic changes in e.g. activity, localization, or interactions (e.g. with DNA, transcription factors, or adaptor proteins) within short time frames. This allows cells to quickly adapt and fine-tune their functions in response to various signals, which is essential for complex processes like brain development [9, 10]. Therefore, understanding the molecular mechanisms underlying the development of CBOs is crucial for deciphering early human brain development and for validating the model for its capability of identifying therapeutic targets for neurological disorders.

Recent studies have elucidated the cellular composition and differentiation trajectories of CBOs through gene expression profiling and assessment of chromatin accessibility [14-21]. Despite these advances, the understanding of how protein dynamics, particularly through PTMs, influence organoid development remains limited. To address this, we generated a first-of-its-kind high-resolution temporal overview of the abundance of proteins and various PTMs in CBOs spanning 28 time points from day 17 to day 200. We focused on PTMs involved in intracellular signaling and neuronal development (including selected active amino acids), such as phosphorylation, various forms of cysteines, lysine acetylation, and sialylated N-glycosylation [22-31]. Using quantitative tandem mass tag (TMT)-based proteomics, PTMomics, and metabolomics, we identified and quantified over 9,300 proteins and various PTM-containing peptides, covering more than 60% of the human proteome. Significant temporal changes in protein abundance and dynamic protein modifications were observed, representing different stages of neurodevelopment. Notably, pathways related to neuronal differentiation and energy metabolism showed changes in both protein abundance and dynamic protein modifications, while calcium signaling was primarily regulated by dynamic protein modifications. We detected a substantial increase in synaptic protein content which correlated with neurotransmitter levels. Astrocytes showed increased abundances beyond day 100 with higher functional properties at later stages. Comparative proteomic analysis revealed similarities between the proteomes of CBOs and human fetal brain tissue, emphasizing the relevance of organoids as models for studying early human brain development. Overall, we believe this temporal proteomic/PTMomic atlas will be a valuable resource for the CBO field.

## Results

### Temporal expression profiles of proteins, PTMs and metabolites in cerebral organoids

To investigate the characteristics of human brain development in CBOs from a protein-centric perspective, we analyzed the expression of proteins, different post-translationally modified peptides (phosphorylation, cysteine modifications, lysine acetylation and sialylated N-glycosylation), and metabolites using mass spectrometry at 28 different time points from day 17-200 (Figure 1A), reflecting different stages of organoid development. The time points included cover periods associated with neural progenitor cells and cell fate acquisition to periods of neuronal maturation and glial cell development [19]. We developed a workflow based on the principles of a previous protocol [32] that enabled the analysis of proteins, multiple PTMs, and metabolites from the same organoid sample at each of the 28 time points (Figure S2), and applied it on CBOs from two different cell lines each subjected to multiple independent differentiations. We identified and quantified 9,395 proteins (Figure 1B), and 38,370 peptides with reversibly modified cysteine residues (Figure 1C), which reflects mainly cysteine residues present in disulfide bonds, but also oxidative modifications. We further identified and quantified 1,775 peptides with sialylated N-glycosylation (Figure 1D), a PTM that is added in the secretory pathway and plays key roles in protein folding, cell-cell interactions, and neurodevelopment [22-24, 33]. In addition, we quantified 29,679 phosphorylated peptides (Figure 1E), which is one of the most common PTMs that regulate several intracellular signaling pathways [25-27]. We identified and quantified 17,012 peptides with a free cysteine residue (Figure 1F), in which the thiol group directly or indirectly can regulate protein activity [28], and 1,154 peptides containing lysine acetylation (Figure 1G), which is mostly known for its roles in gene regulation and cellular metabolism [29-31]. We furthermore identified and quantified 396 metabolites in the organoid samples (Figure 1H). Together, the proteins and modified/cysteine-containing peptides represent a total of 12,346 proteins, covering more than 60% of the human proteome [11, 34]. We observed that in most cases, more than 40% of the identifications in a dataset showed significant temporal expression changes during organoid development (Figure 1B-H).

**Figure 1:**
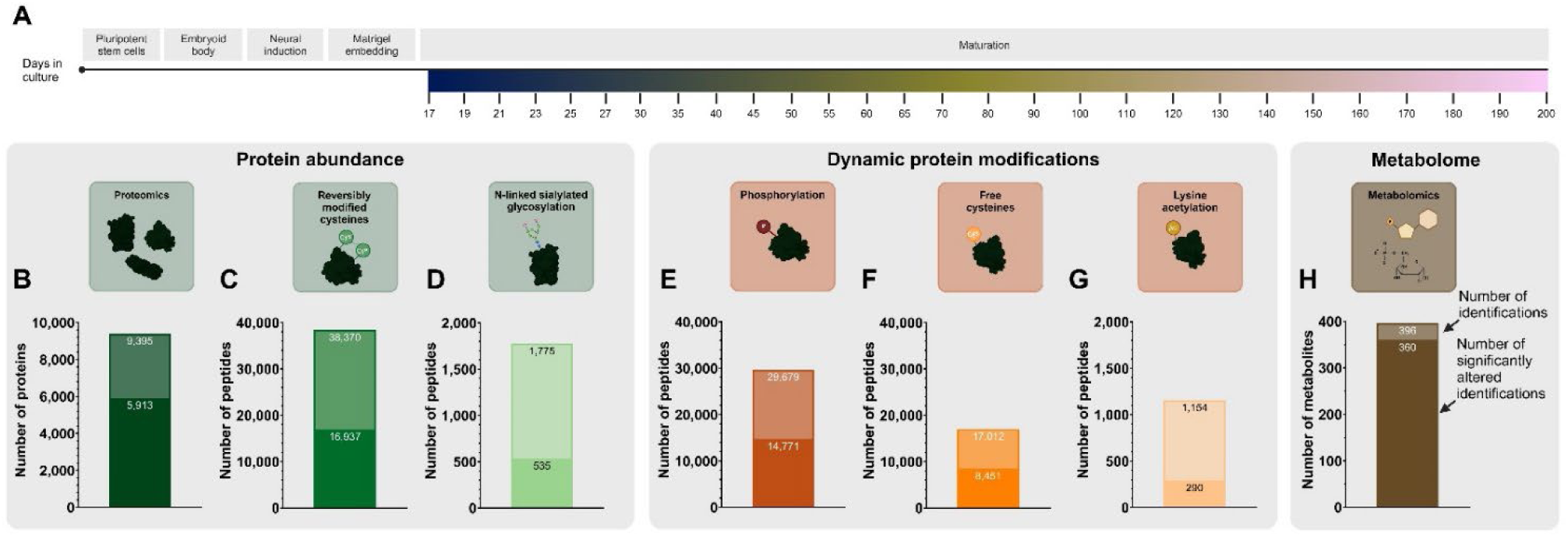
Protein-centric multi-omic high-resolution temporal atlas of cerebral organoid development. **A**: Schematic overview of the experimental setup for culturing cerebral organoids and time points included in the study. **B-H**: Overview of the number of identified proteins (**B**), post-translationally modified peptides (**C-G**), and metabolites (**H**).

Since some of the PTMs included in this study have roles in protein folding and stabilization while others have roles in cellular signaling, we chose to group the proteomics and PTM-specific proteomics (PTMomics) data into two categories: reflectors of protein abundance (proteomics, reversibly modified cysteines, and N-linked sialylated glycosylation), and reflectors of dynamic protein modifications (phosphorylation, free cysteines, and lysine acetylation). Principal component analysis revealed that all datasets showed separation of organoid samples based on organoid age at the time of analysis (Figure 2A-C+G-I+M) with a positive association between one of the first two principal components and organoid age (Figure S4), showing that gradual progression of organoid development is reflected in the abundance of proteins, dynamic protein modifications, and metabolites. Organoids generally showed less variability at early time points, whereas intermediate and later time points showed more variability in most datasets (Figure 2A-C+G-I+M), which could be related to differences between cell lines (Figure S3+Figure S4), likely due to different cellular compositions caused by differences in differentiation trajectories. Protein abundance reflectors that had the highest contributions to the first principal component included proteins involved in regulation of chromatin (SMARCA4, ZMYM2, TLK2, NSD2, SUPT16H, SMARCA5, KAT5, and CREG1), and cellular morphology and cytoskeletal organization (ZMYM4, SLC2A13, SCARF2, LAMA5, and CD63). Among the top contributors to the second principal component were proteins related to neuronal axons, synapses, vesicular transport, and calcium homeostasis (NAPB, PPFIA3, EVI5L, WDR47, ATP6V1G2, DLG4, NRXN2, BRSK1, PALM, RNF157, AAK1, CACNB1, MAP1B, NCAM2, VSIG10, OLFM1, CACNA2D1, SSR2, CADM2, and ASPH) (Figure 2D-F). Among the reflectors of dynamic protein modifications, phosphorylation on proteins involved in transcriptional regulation (PHC3, ARMC10, ZNF532, BRD3, MBD3, JAZF1, and PHF6) and synapse function, ion transport and microtubule organization (EPB41L1, SHANK2, CTNNA3, SPTBN1, LRP1, CEP170, and SLCO3A1) contributed the most to the first two principal components (Figure 2J). Among the proteins identified with a free cysteine residue, proteins involved in gene regulation and protein synthesis and folding (PPIH, RNF213, EIF2S2, TATDN1, NAA15, RAN, and KPNB1) contributed the most to the first principal component, whereas SLC2A13, a protein found in neuronal growth cones and axons, as well as proteins involved in mitochondrial functions related to redox homeostasis and the citric acid cycle (ACO2, IDH3A, PRDX5, and RNH1) contributed the most to the second principal component (Figure 2K). Of the proteins with an acetylated lysine, proteins involved in cellular metabolism (GOT2, IDH2, GAPDH, COX4I1, and MDH1) were among the top contributors to the first principal component, whereas proteins involved in gene regulation (ZFR, RNPS1, TRIM28, BAZ1B, H2BC18, and H2AJ) were among the top contributors to the second principal component, and proteins involved in calcium homeostasis (ANXA6, CPNE1, VAPB, and CACYBP) contributed to both the first and second principal component (Figure 2L). For the metabolomics data, the top contributors to the first and second principal components included regulators of amino acid metabolism and amino acid precursors for neurotransmitters (e.g. tryptophan) (Figure 2N). Overall, this data provides a thorough description of the key protein abundance and dynamic modification changes occurring during CBO development.

**Figure 2:**
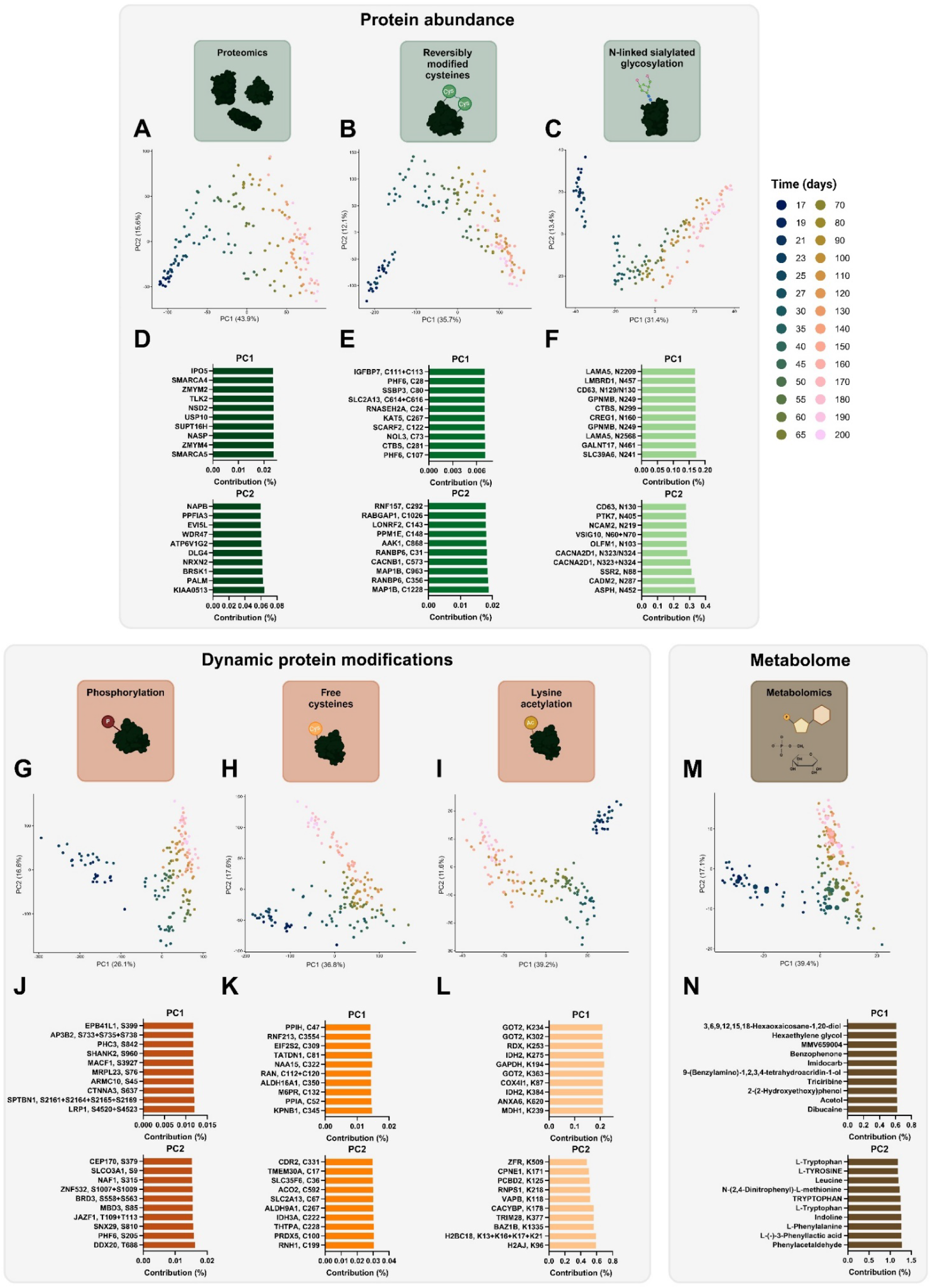
Cerebral organoids show temporal abundance profiles for proteins, post-translational modifications, and metabolites during organoid maturation. **A-C**: Principal component analysis of proteomics (**A**), reversibly modified cysteines (**B**), and N-linked sialylated glycosylation (**C**) data, which reflect temporal profiles of protein abundance. **D-F**: The 10 features (proteins/peptides) that contribute the most to the first and second principal component for the proteomics (**D**), reversibly modified cysteines (**E**), and N-linked sialylated glycosylation (**F**) data. **G-I**: Principal component analysis of phosphorylation (**G**), free cysteines (**H**), and lysine acetylation (**I**) data, which reflect temporal profiles of dynamic protein modifications. **J-L**: The 10 features (peptides) that contribute the most to the first and second principal component for the phosphorylation (**J**), free cysteines (**K**), and lysine acetylation (**L**) data. **M**: Principal component analysis of metabolomics data. **N**: The 10 features (metabolites) that contribute the most to the first and second principal component of the metabolomics data.

### Neuronal development and energy metabolism proteins show temporal changes in both abundance and dynamic modifications

We subjected the protein abundance and dynamic protein modification reflectors that showed altered abundance across organoid development to unsupervised clustering analysis. This resulted in grouping of protein abundance reflectors into 9 clusters (Figure 3A+B). Some clusters contained proteins with the highest abundance at early time points (e.g. cluster 1, 3, 5, and 9) (Figure 3A). These clusters were enriched in proteins involved in cell proliferation, Notch signaling, chromatin remodeling, and RNA splicing (Figure 3C), all related to progenitor cell function. Other protein abundance clusters contained proteins whose abundance increased during organoid development (e.g. cluster 2, 4, and 7) (Figure 3A). Cluster 2 contained proteins enriched in KEGG pathway terms related to metabolism (carbon metabolism and oxidative phosphorylation) and neuron maturation and synaptic function (axon guidance, synaptic vesicle cycle, retrograde endocannabinoid signaling, ECM-receptor interaction, and glutamatergic synapse) (Figure 3C). Proteins in cluster 4 and 7 were enriched in terms related to metabolism (carbon metabolism in cluster 4 and carbon metabolism, oxidative phosphorylation, and fatty acid degradation in cluster 7), ECM-receptor interactions, phosphatidylinositol signaling, and lysosomes (also reflected in a higher proportion of features in cluster 7 arising from sialylated N-linked glycopeptides) (Figure 3B+C). Cluster 6 also contained proteins enriched in KEGG terms related to neuronal development, but in contrast to cluster 2, 4, and 7, showed the highest abundance at early time points (Figure 3C). Interestingly, cluster 8 contained proteins with the highest abundance around day 30-80, and was enriched in axon guidance, glutamatergic synapse, and motor proteins terms (Figure 3C).

**Figure 3:**
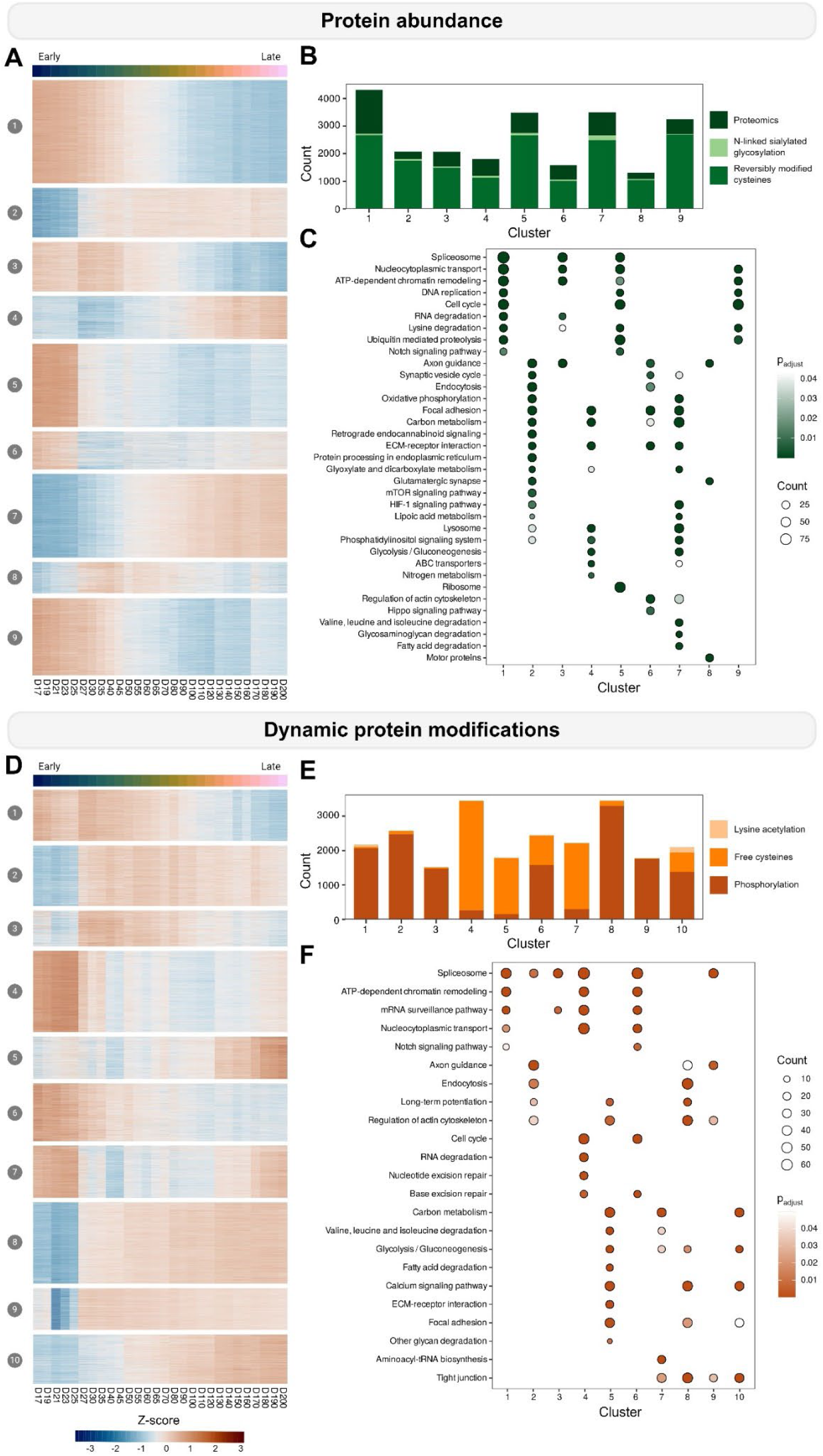
Temporal profiles of protein abundance and dynamic protein modification reflectors. **A**: Heatmap of clusters of protein abundance reflectors and their relative expression temporal profiles across cerebral organoid development. **B**: Overview of the number of cluster members in each protein abundance cluster and the proportion of members from the proteomics (dark green), reversibly modified cysteines (green), and N-linked sialylated glycosylation data (light green), respectively. **C**: KEGG pathway enrichment results for protein abundance clusters. Circle size reflects the number of proteins identified in a protein abundance cluster, whereas circle color reflects enrichment p-value after Benjamini-Hochberg adjustment for multiple testing. **D**: Heatmap of clusters of dynamic protein modification reflectors and their relative temporal profiles across cerebral organoid development. **E**: Overview of the number of cluster members in each dynamic protein modification cluster and the proportion of members from the phosphorylation (dark orange), free cysteines (orange), and lysine acetylation data (light orange), respectively. **F**: KEGG pathway enrichment results for dynamic protein modification clusters. Circle size reflects the number of proteins identified in a dynamic protein modification cluster, whereas circle color reflects enrichment p-value after Benjamini-Hochberg adjustment for multiple testing.

The dynamic protein modification reflectors were grouped in 10 clusters (Figure 3D+E). Like the protein abundance clusters, we detected clusters of proteins with dynamic modifications related to progenitor cell functions such as proteins involved in chromatin remodeling and RNA splicing in cluster 1, 4, and 6, as well as proteins related to cell proliferation in cluster 4 (Figure 3F). These clusters showed the highest abundance at early time points and decreased over time (Figure 3D). In clusters where the abundance of modified peptides increased over time (such as cluster 2, 8, 9, and 10), proteins were mainly modified by phosphorylation (Figure 3E). Modified proteins detected in cluster 2 and 9 were enriched in terms related to neuronal development, synaptic function, and RNA splicing, and showed the highest abundance around D60-80 (Figure 3F + Figure S7). The enrichment of neuronal terms was also reflected in enrichment of cellular components related to axons and growth cones in these clusters (Figure S8B). Cluster 8 and 10 showed enrichment of tight junction and calcium signaling pathways and the proteins carrying modifications showed increased abundance over time (Figure 3F + Figure S7). Cluster 5 also contained modifications on proteins involved in calcium signaling in which the abundance increased after D100-120 (Figure 3D+F). Apart from enrichment of calcium signaling, modified proteins identified in cluster 5 were also related to metabolic and adhesion pathways and enriched in cellular components related to cell-ECM interactions and postsynaptic density (Figure 3F + Figure S8B). Metabolic pathway terms enriched were related to carbon metabolism, amino acid metabolism, and fatty acid degradation, which is highly abundant in astroglia and could be related to their presence in the organoids. Another dynamic modification cluster of primarily phosphorylated proteins had increased abundance at earlier time points and decreased after D70-80 (cluster 3; Figure 3D+E). These proteins were enriched in splicing-related processes and cellular components related to neuronal structures such as axons and growth cones (Figure 3F + Figure S8B).

Together, this shows that the protein expression profiles of CBOs replicate the timing of processes involved in early stages of human brain development. This includes similarities with previous transcriptomic-based studies of neural organoids [19, 20]. Our data also shows that PTMs on proteins involved in processes related to neuronal maturation and energy metabolism show altered abundance during organoid development, indicating that these processes likely are regulated by dynamic protein modifications. Free cysteines were highly abundant at early and late time points, whereas protein phosphorylation was abundant on proteins involved in both progenitor functions as well as in proteins related to neuronal maturation, suggesting that protein phosphorylation have a variety of functions during organoid development.

### Axon development is promoted by increasing protein abundance and dynamic modifications

We examined the clusters related to axonal guidance for both protein abundance and dynamic protein modification reflectors. In both protein abundance and dynamic protein modification clusters, we observed guidance molecules and their receptors as well as adhesion molecules, G-protein effectors, and protein kinases (Figure 4A+B). Examples include semaphorins, plexins, Robos, Wnts, BMPs, Eph receptors, Slits, Uncs, CDC42, Racs, and protein kinases such as CDK5, PAK family kinases, FYN, PI3K, SRC, CAMK2 kinases, and GSK3B. The modified axon guidance proteins identified in dynamic protein modification clusters 2 and 9 were mostly phosphorylated (Figure 4B). Some axon guidance proteins were present in protein abundance clusters with the highest abundance around D30-40 or earlier (e.g. BMPR2, EFNB2, SEMA5B, SLIT3, UNC5B, and WNT4). Others were present at higher abundance from D40-50 and onwards (e.g. CAMK2B, CDC42, PIK3CA, PLXNB2, PRKCA, RAC1, and SEMA4G) (Figure 3A + Figure 4A). Modifications on some of these proteins, primarily by phosphorylation, were more abundant at slightly later time points (around D60-80 and onwards) (Figure 3D). Most of the axon guidance-related proteins, which we detected as phosphorylated, were receptors, protein kinases, or regulators of cytoskeleton (Figure 4B). Phosphorylation sites on proteins, where the phosphorylation of these sites has previously been shown to regulate actin cytoskeleton dynamics include pS937 on SSH1 [35], pY68 on CFL1 [36], pS181 on PAK4 [37], pS144 on PAK1 [38], and pT514 on DPYSL2 [39, 40]. SSH1 and PAK4, regulates CFL1-mediated disassembly of actin filaments directly (SSH1) or indirectly (PAK4) [41, 42]. SSH1 phosphorylation of S937 reduces its activity [35], whereas CFL1 phosphorylation of Y68 has been associated with CFL1 degradation [36], indicating that actin cytoskeleton could be stabilized over time as would be required for growth of neuronal processes. We further identified activating phosphorylation sites on the Src family non-receptor kinases SRC (pS17) [43] and FYN (pS21) [44] as well as on PAK1 (pS144) [38], PAK2 (pS141) [45], and MAPK3/ERK1 (pT202) [46] (Figure 4B), suggesting that these kinases had higher activities as the organoids matured. SRC, FYN, PAK1, PAK2, and MAPK3 were central nodes in the protein-protein interaction network, displaying interactions with guidance molecule receptors. These have roles in multiple aspects of neuronal development and maturation such as neuronal migration, axon growth and synaptogenesis, indicating that these findings may have dual roles and not be specifically related to axon guidance. Related to this we observed an increased abundance of motor proteins of the kinesin, dynein, and myosin superfamilies around D30-80 (Figure 4C), which were associated with neuronal axons and synapses (Figure S8A). Motor proteins are related to neuronal polarity, and by transporting cargoes to and from the soma, they mediate intracellular transport in neurons [47]. For example, KIF1A, KIF1B, and KIF5C transport synaptic vesicle precursors, presynaptic membrane components, mitochondria, and mRNA for local protein synthesis [48-52]. Delivery of these components to the neuronal axons is critical for proper maturation and formation of synapses since their localization can regulate their functions [47]. At periods partially overlapping with the presence of axonal guidance and motor proteins and the modifications on axonal guidance proteins, we identified dynamically modified proteins related to calcium signaling. This was interesting since calcium signaling is required for synaptic signaling and has also been implicated in neurite growth [53, 54]. In contrast to axonal guidance proteins, proteins involved in calcium signaling were modified both by protein phosphorylation and had abundant free cysteine residues (Figure 4D), which could regulate activity and interactions. They were related to growth factor and neurotransmitter receptors (e.g. for EGF, glutamate, and serotonin), phospholipase C, protein kinase C, calcium channels (e.g. IP3-related and voltage-dependent), calmodulin, and CAMK2 kinases. Together, these intracellular signaling proteins regulate the cytoplasmic levels of calcium, and their PTMs could affect local intracellular calcium fluctuations [55]. Phosphorylation of PLCB3 at S537 and EGFR at S1166 can both be catalyzed by CAMK2A [56, 57], and pS537 of PLCB3 has previously been associated with increased cellular motility [58]. On CAMK2B and CAMK2D, we identified peptides carrying both a free cysteine residue (C290) and phosphorylation at T287, which is known as the autophosphorylation site of CAMK2 kinases [59, 60], indicating that they might be active at later stages. Phosphorylation of PRKCA at S226 induces its kinase activity [61] and could participate in calcium mobilization. We furthermore observed regulatory phosphorylation sites (pS257 and pS521) on STIM1 [62], a protein which has been linked to calcium dynamics in growth cones [63], in dynamic protein modification clusters 8 and 10. STIM2, which has been associated with dendrites of more mature neurons [64], was also modified by both phosphorylation and had increased abundance of free cysteine residues at higher levels above D130, however, the function of these modifications is not known. Together, these results indicate that axon development is promoted by increasing both protein abundance and dynamic modifications, whereas calcium signaling is mainly dynamically altered during CBO development.

**Figure 4:**
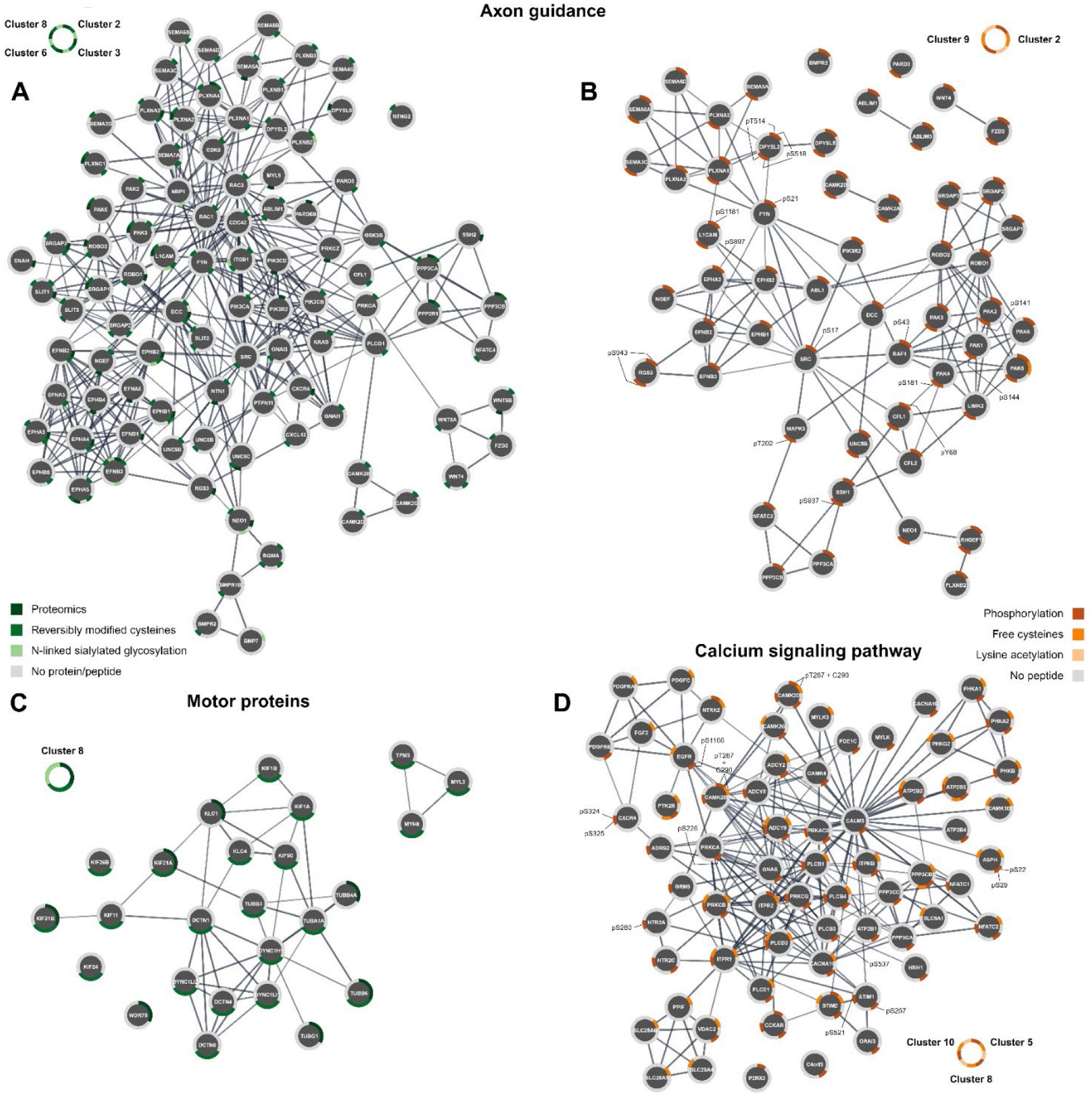
KEGG pathway terms related to neuronal development and function. **A-B:** STRING protein-protein interaction diagrams of proteins related to enrichment of axon guidance in protein abundance clusters (**A**) and dynamic protein modification clusters (**B**). **C:** STRING protein-protein interaction diagram of motor proteins from the protein abundance cluster 8. **D:** STRING protein-protein interaction diagram of proteins from the dynamic protein modification clusters involved in the calcium signaling pathway. Protein-protein interactions with high confidence (>0.7) are displayed. Line thickness indicate confidence of protein-protein interaction (increasing thickness = higher confidence). Circles show the gene name of proteins. The outer ring in **A** and **C** denotes the presence of protein/peptide features from protein abundance datasets in the given cluster. The outer ring in **B** and **D** denotes the presence of peptide features from dynamic protein modification datasets in the given cluster.

### Increasing synaptic protein content correlates with neurotransmitter levels

As our analysis had identified clusters of proteins related to neuronal maturation with increasing temporal abundance and the involvement of dynamic protein modifications in related processes, we asked whether the abundance profiles of proteins related to synapses were similar. We selected proteins present in the presynapse and in synaptic vesicles, as well as proteins present in the postsynapse, including neurotransmitter receptors, based on information in the Gene Ontology knowledgebase [65, 66]. We observed that many synapse-related proteins, including SNAP25, SYP, SV2A, SYN1, SHANK3, and GRID2, showed the highest relative abundance around day 60-80 (Figure 5A-D). Others showed higher abundances above day 100, e.g. VAMP1, HOMER1, SIGMAR1, and GRIK3 (Figure 5B-D). However, some scaffold proteins that localize to synapses, such as BSN, PCLO, and SCRIB, had a relative decrease in abundance over time (Figure 5A+C), which could be related to an expansion of the cellular volume as neurons extend axonal and dendritic processes. Using metabolomics, we quantified the intracellular levels of neurotransmitters and their metabolic precursors or intermediates. We found that excitatory glutamate levels increased and peaked around day 100, whereas the levels of the glutamate precursor glutamine remained at high levels from day 100-200 (Figure 5E) and the inhibitory neurotransmitter GABA was not detected. We detected N-acetylaspartylglutamate (NAAG), an abundant dipeptide neurotransmitter in the CNS, at the highest levels around D60-80 (Figure 5E) correlating well with the alterations in axon guidance proteins. The non-classical neurotransmitter taurine had increased abundance until day 80-100, and precursors for other neurotransmitters such as serotonin and acetylcholine had the highest abundance at late time points (Figure 5E). To confirm the temporal presence of mature neuronal and synaptic proteins, we performed Western blot analysis for MAP2 (Figure 5F + Figure S9) and SYN1 (Figure 5G + Figure S9). Corresponding to our proteomic analysis, the relative abundances of MAP2 and SYN1 peaked around day 65-100 (Figure 5F+G). We used immunohistochemistry to evaluate the localization of SYN1 at day 40, 80, and 170 (Figure 5H). At day 40, SYN1 showed close nuclear localization and sparce expression of TUBB (Figure 5H), indicating limited neuronal differentiation. SYN1 localization was more distal at day 80, whereas CBOs at day 170 showed increased abundance of SYN1 and TUBB-positive filaments (Figure 5H), indicative of axon outgrowth and more mature neurons at late time points. Since synapses require the right spatial localization of synaptic components, including proteins, as well as loading of neurotransmitters into synaptic vesicles, and since some neurotransmitters such as glutamate can also have metabolic functions, we used a multi-electrode array to evaluate the functional properties of neurons at day 120, 140, 160, 180, and 200. We detected spontaneous activity at all time points, and the mean firing rate was significantly increased at day 180 and day 200 (Figure 5I), indicating that neurons matured into functional cells and displayed increased levels of synaptic activity at late time points.

**Figure 5:**
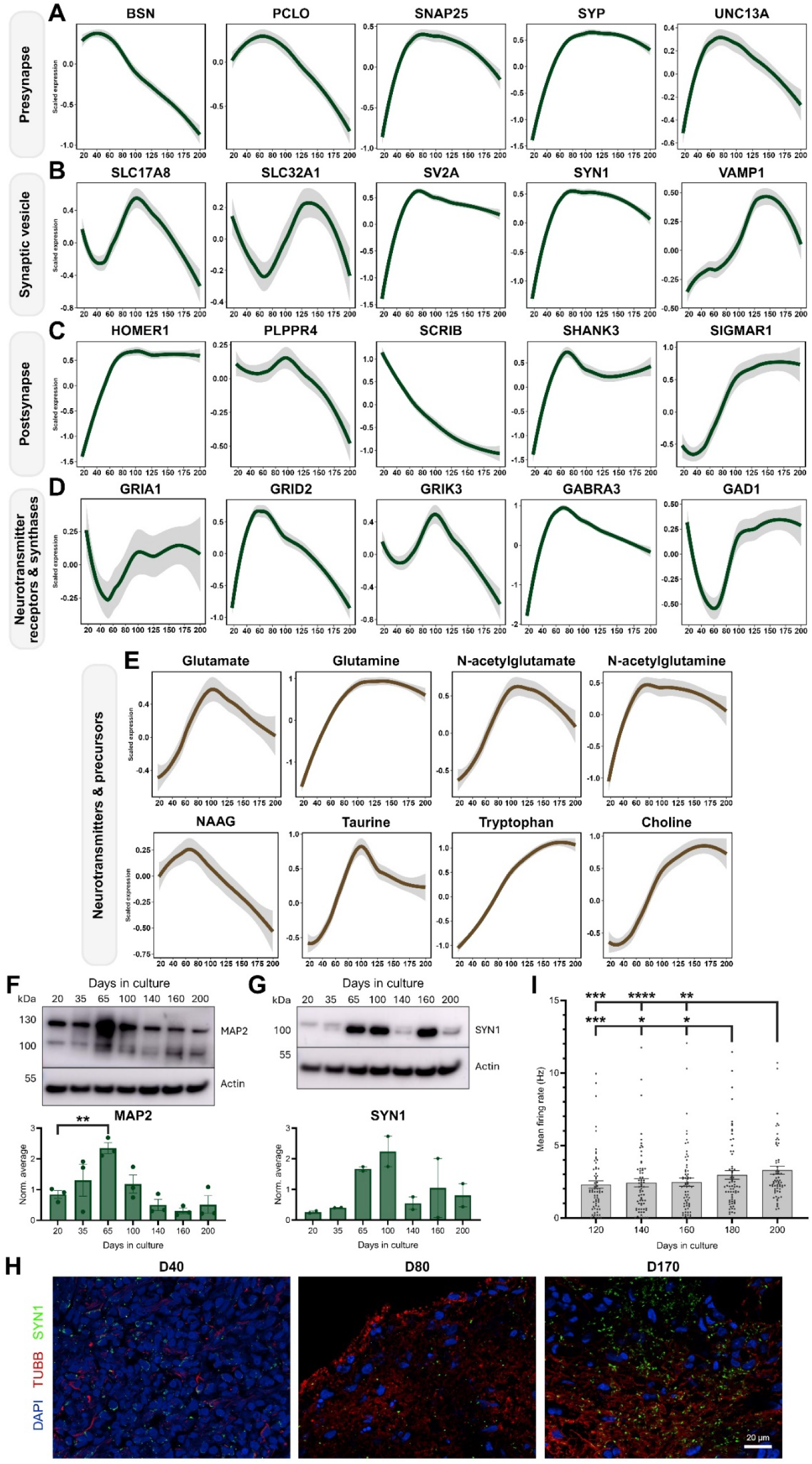
Abundance of synaptic proteins and neurotransmitters during cerebral organoid development. **A-D:** Average z-scaled abundance profiles of proteins from the presynapse (**A**), synaptic vesicles (**B**), postsynapse (**C**), and neurotransmitter receptors/synthases (**D**). **E:** Average z-scaled abundance profiles of neurotransmitter molecules and related metabolites. NAAG: N-acetylaspartylglutamate. The average z-scaled abundances in **A-E** were fitted to a LOESS (locally estimated scatterplot smoothing) regression model. **F-G:** Representative western blots and quantification of MAP2 (**F**) and SYN1 (**G**) levels in cerebral organoids at day 20, 35, 65, 100, 140, 160, and 200. Protein levels were normalized to β-actin and the average of all samples in each blot. Mean ± SEM. ** p<0.01 (One-way ANOVA). **H:** Immunohistochemistry for DAPI (blue), beta tubulin (TUBB, red), and SYN1 (green) in cerebral organoids at day 40, 80, and 170. Scalebar: 20 μm. **H:** Multi-electrode array recording of spontaneous neuronal activity in 10 organoids at day 120, 140, 160, 180, and 200. Mean ± SEM. Points denote individual electrodes. * p<0.05, ** p<0.01, *** p<0.001, **** p<0.0001.

### Astrocytes are present beyond day 100 and correlated with fatty acid degradation

Apart from the clusters related to neuronal differentiation, the clustering analysis showed the presence of protein abundance and dynamic protein modification clusters related to metabolism (Figure 3A-F). These clusters had increased abundances over time and included proteins related to glycolysis and oxidative phosphorylation, which is likely linked to increased levels of neurons and synapses. However, we also observed increased abundance of proteins related to fatty acid β-oxidation over time. Since astrocytes have high metabolic activities and are the predominant cell type performing fatty acid β-oxidation in the brain, this led us to investigate whether these findings might be related to the presence of astrocytes in the CBOs. We observed that TMSB15A, a marker of early astrocytes, peaked at the earliest time points and decreased over time (Figure 6A). Astrocytic markers that describe more mature astrocytes such as GFAP, GLAST and S100B increased in abundance over time and reached a plateau around day 100. The astrocytic marker AQP4 had a shift in abundance upon day 100 and showed increased protein levels from day 100 to 200 without reaching a plateau (Figure 6A). This suggests the presence of astrocytes in cerebral organoids after day 100. Markers associated with oligodendrocytes or oligodendrocyte precursor cells mostly decreased in abundance over time and OLIG1 and OLIG2 were not detected (Figure S10), suggesting that oligodendrocytes were likely not present in the CBOs. Western blot analysis confirmed that the abundance of GFAP was significantly increased at day 100 and 140 compared to day 20 (Figure 6B + Figure S9). Immunohistochemistry showed higher abundance of GFAP in CBOs at day 80 and 170 compared to day 40 with localization at the edge of the organoid as well as between MAP2-positive neurons (Figure 6B + Figure S11). Notably, by day 170, GFAP-positive cells surrounding MAP2-positive neurons exhibited a more ramified morphology (Figure 6B), suggesting more advanced astrocytic differentiation within the CBOs.

**Figure 6:**
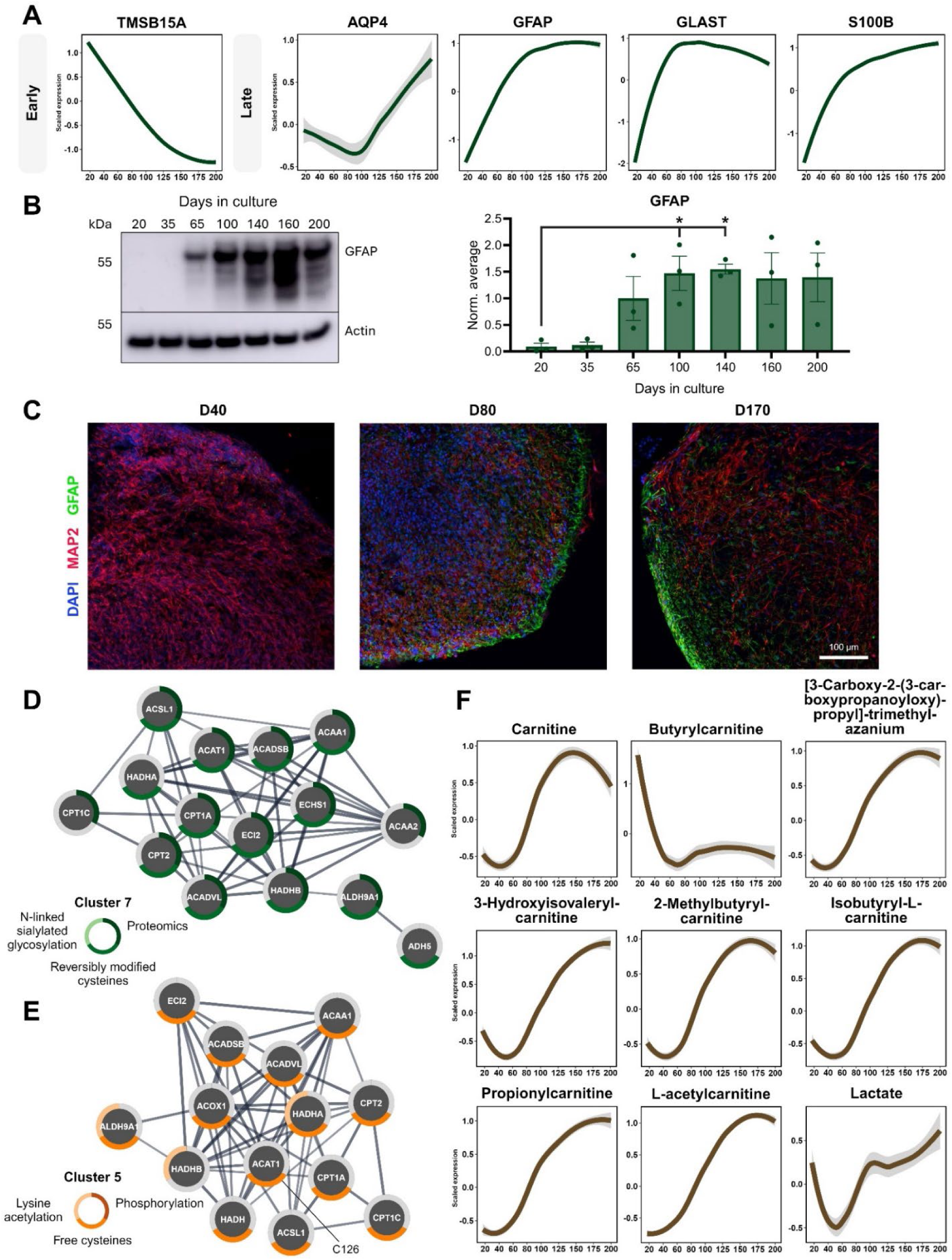
Abundance of astroglial proteins above day 100 in cerebral organoids correlate with me-tabolic changes related to fatty acid oxidation. **A:** Average z-scaled abundance profiles of astroglial markers. **B:** Representative western blots and quantification of GFAP levels in cerebral organoids at day 20, 35, 65, 100, 140, 160, and 200. Protein levels were normalized to β-actin and the average of all samples in each blot. Mean ± SEM. * p<0.05 (One-way ANOVA). **C:** Immunohistochemistry for DAPI (blue), MAP2 (red), and GFAP (green) in cerebral organoids at day 40, 80, and 170. Scalebar: 100 μm. **D-E:** STRING protein-protein interaction diagrams of proteins from protein abundance cluster 7 (**D**) and dynamic protein modification cluster 5 (**E**) related to fatty acid degradation. Protein-protein interactions with high confidence (>0.7) are displayed. Line thickness indicate confidence of protein-protein interaction (increasing thickness = higher confidence). Circles show the gene name of proteins. The outer ring denotes the presence of protein/peptide features from protein abundance datasets (**D**) or peptide features from dynamic protein modification datasets (**E**) in the given cluster. **F:** Average z-scaled abundance profiles of carnitine, acylcarnitines, and lactate. The average z-scaled abundances in **A** and **F** were fitted to a LOESS (locally estimated scatterplot smoothing) regression model.

Some of the fatty acid degradation proteins we identified in protein abundance and dynamic protein modification clusters have been described as astrocyte markers, e.g. CPT1A, CPT2, ECI2, and HADHA (Figure 6D+E). We observed that the modified fatty acid degradation proteins were mostly identified with a free cysteine residue (Figure 6E). The modification abundance on these proteins showed elevation after increased abundance of the unmodified proteins (Figure 3A+D), indicating that the modification of these proteins could affect their enzymatic functions. CPT1A and CPT2 are involved in the carnitine-mediated transport of fatty acids to mitochondria for their subsequent β-oxidation. We observed that ACAT1, which catalyzes the last step of fatty acid oxidation, had higher abundance of a free cysteine in its active site (C126) [67] at late time points (Figure 6E), indicative of increased activation of β-oxidation. Additionally, the levels of carnitine and acylcarnitines increased over time, reaching the highest abundance around day 150 in most cases (Figure 6F), indicating that metabolic activity in astrocytes peaks at late time points in CBOs. Further supporting this was the levels of lactate, a product of glycolysis, which is produced in proliferative cells as well as in astrocytes to support neuronal metabolism. As expected, lactate was abundant at early time points, but also showed increased abundance at later time points above day 60-100 and interestingly, lactate levels increased further in CBOs around day 150-200 (Figure 6F). Together, this could suggest that astrocytes are abundant around day 100 in CBOs but display increasing functional properties at later time points.

### Cerebral organoids show proteomic similarity with human fetal brain tissue

To evaluate the physiological relevance of CBOs with respect to their proteome, we compared the proteome of CBOs to the proteome of human brain tissue described by Djuric et al. in 2017 [68]. Djuric et al. dissected and analyzed fetal brain tissue from the ventricular zone, intermediate zone, subplate, and cortex regions, respectively. We compared the proteomics results from CBOs from day 17 to day 200 to human brain proteomes from gestational week (GW) 16 to 41. We observed that CBOs collected at early time points (day 17-30) correlated mostly with early-midgestational ventricular zone areas and partially with cortex areas (Figure 7A), likely due to neurogenic events. CBOs collected at time points above day 80 correlated with mid-late gestational intermediate zone, subplate and cortex areas (Figure 7A). From around day 35-80, CBOs generally showed lower levels of correlation with fetal brains in most areas, but the correlation between CBOs and fetal brain tissue from the subplate gradually increased from day 55 to 80 (Figure 7A). This indicates that this period could present a transition state within CBOs related to neuronal migration and cortical lamination. Among proteins that showed positive correlation between CBOs and fetal brains were proteins related to cell proliferation, chromatin remodeling, and nucleocytoplasmic transport having highest abundances in the ventricular zone of human brains as well as in early time points of CBOs (Figure 7B). Proteins correlating with intermediate zone and subplate regions from human brains were related to membrane remodeling, intracellular vesicular transport, and migration (Figure 7B). Proteins with the highest abundance in cerebral cortex mostly showed increased abundance in CBOs after day 40-60 and were associated with enzymes related to metabolism and sialylation. These results indicate that the proteome of CBOs shares similarities with that of the *in vivo* developing cerebral cortex at fetal stages.

**Figure 7:**
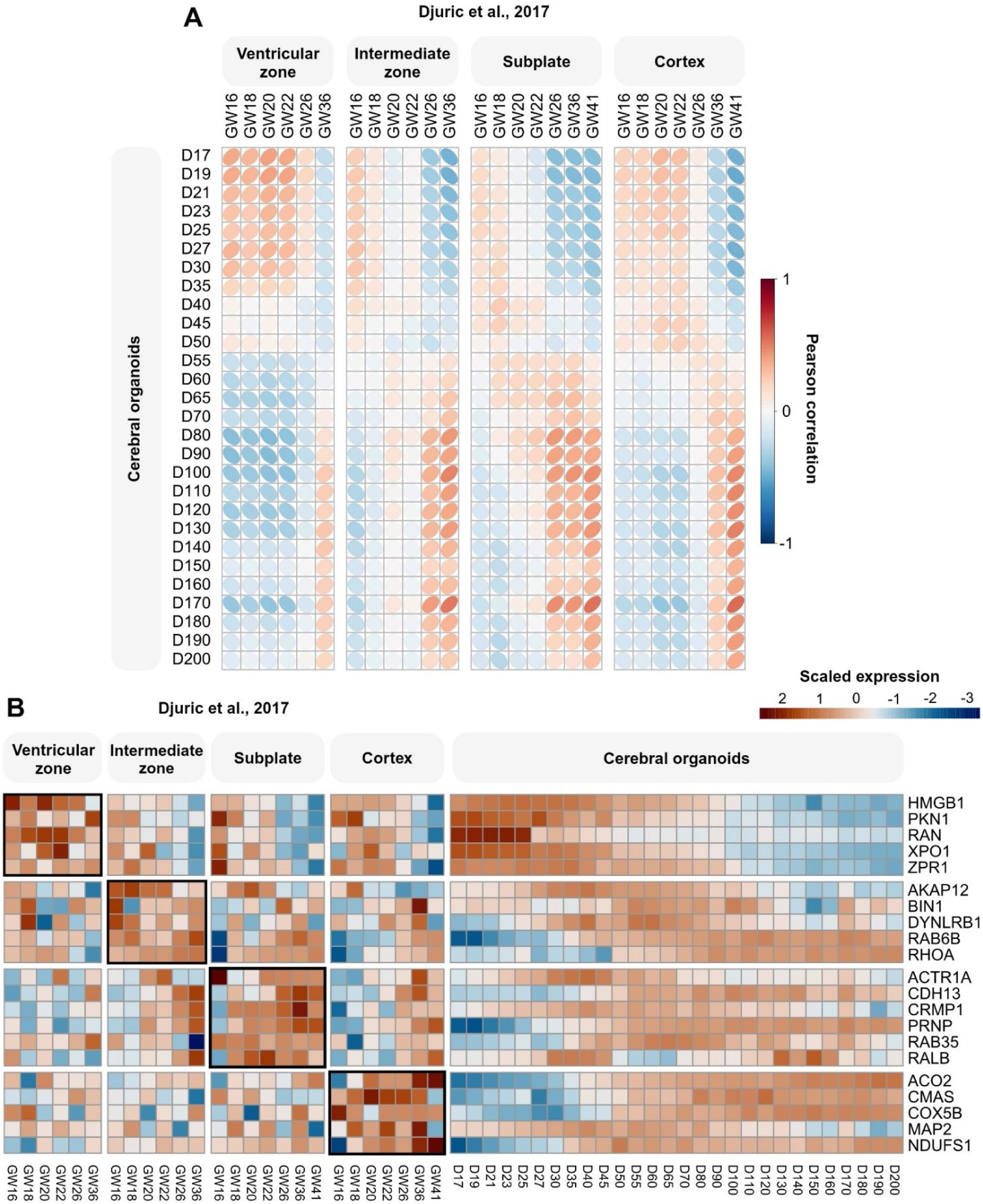
Spatiotemporal comparison of cerebral organoid proteome and fetal brain tissue. **A:** Pearson correlation between cerebral organoid proteomes from day 17-200 and proteomes of ventricular zone, intermediate zone, subplate, and cortex regions dissected from FFPE fetal brain samples from GW16-GW41 by Djuric et al. [68]. **B:** Examples of proteins with similar relative expression profiles between CBOs and fetal brain compartments.

## Discussion

Proteomics and PTMomics hold great potential in neural organoid research, as they complement more widely used transcriptomic and epigenomic approaches. Proteomics and PTMomics analyze the functional outcomes of epigenetic modifications and alterations in gene expression by measuring the dynamics of proteins and PTMs. This can aid in elucidating new aspects of signaling mechanisms involved in early human brain development and evaluate the applicability of CBOs as a model of the human fetal brain. With this study, we present a comprehensive temporal atlas of proteomic and PTMomic changes during the development of CBOs across 28 time points from day 17 to day 200. By integrating proteomic and PTMomic data, we covered more than 60% of the human proteome and showed that protein abundances and dynamic protein modifications display significant temporal changes during CBO development related to different stages of neurodevelopment. We demonstrate that certain pathways were associated with changes in protein abundance and dynamic protein modification abundance, such as pathways involved in neuronal differentiation and energy metabolism, whereas pathways related to calcium signaling were mostly associated with changes in dynamic protein modification abundance. These findings highlight the role of PTMs in regulating protein functions in response to developmental cues in key developmental processes and physiological functions that are critical for proper brain function.

We found that proteins related to axon guidance were mostly modified by protein phosphorylation. This included guidance receptors and protein kinases that regulate cytoskeleton dynamics (PAK1, PAK2, MAPK3, SRC, and FYN). Axon guidance receptors and the dynamics of actin filaments are involved in multiple aspects of neuronal differentiation, including regulation of neuronal fate, neuronal migration, axon guidance and synaptogenesis [69, 70]. Thus, the phosphorylation of these proteins could alter one or more of these processes. We observed increased abundance of active phosphorylated PAK1 and PAK2 kinases from day 27 and onwards. PAKs are downstream effectors of Rho family GTPases [71, 72], play important roles in neuronal morphology [71-73], neuronal migration [74, 75], and axon development [76, 77], and have been linked to neurodevelopmental diseases [75, 78-81]. Active phosphorylated SRC kinases such as Src and Fyn, which we observed peaked in CBOs around day 50-100, have been proposed to be essential for netrin-mediated axon outgrowth [82] and can regulate the activity of PAKs in the filopodia of growth cones [83]. Thus, we show that there is a temporal aspect of axonal guidance protein regulation in CBOs, but further investigation of spatial aspects such as protein localization as well as analysis of substrates of Src and PAK kinases could further help to understand their role in neuronal differentiation.

Partially overlapping with the temporal profiles of axon guidance proteins, we observed increasing abundances of synaptic proteins and neurotransmitters around day 60-100. However, neurons displayed significant changes in synaptic firing rates only at later time points, suggesting that changes in subcellular protein localization likely precede synaptic activity. Another possibility is that synaptic proteins can have non-synaptic functions in less mature neurons before synaptogenesis. Multiple synaptic proteins have been identified in growth cones (e.g. PCLO, SNAP25, SV2A, and Vamp) [84-86] and could therefore also participate in axon outgrowth-related membrane trafficking at earlier time points. We observed that the relative levels of neuronal and synaptic proteins decreased at later time points, where other findings suggest maturation of neurons. There can be multiple explanations for this. One could be the cellular composition of CBOs, which changes over time and is reflected in the average abundance of proteins across organoids. The increase in proteins of non-neural cell types such as glial cells will result in a relative decrease in other proteins such as neuronal proteins. Another potential explanation could be related to the morphology of neurons, which changes as they differentiate and extend projections. The extension of neuronal projections expands the neuronal surface area. If this is not accompanied by similar increases in the level of synaptic proteins, their relative abundance levels will decrease.

Interestingly, we found that calcium signaling proteins were mainly altered by dynamic protein modifications such as phosphorylation and the presence of a free cysteine residue. The abundance of these modifications was higher at later time points in CBOs and was related to both neurotransmitter receptors, ion channels and protein kinases such as CAMK2B and CAMK2D. The roles of calcium signaling in neural tissue are diverse. In neurons, calcium-related signaling can regulate neuronal subtype specification, outgrowth of neurites and axons and synaptic transmission [87]. However, glial cells such as astrocytes also express neurotransmitter receptors and calcium channels, which are involved in the regulation of release of gliotransmitters, synaptic transmission and neuronal spiking [88, 89]. Thus, our findings could have several implications depending on whether the changes we observed are related to neurons, glial cells, or both. CAMK2 kinases, in which we observed higher abundance of autophosphorylation with concurrent presence of a free cysteine in its vicinity at late time points, have been associated with neurodevelopmental disorders [90, 91], in which rare genetic variants altered the autophosphorylation and hereby activity of CAMK2D [90]. Our findings highlight that dynamic protein modifications play an important role in calcium signaling pathways and further investigation into the functional effects of these alterations could provide more insight into their mechanistic roles in neurodevelopment.

We further detected astrocytes in CBOs beyond day 100 with increased abundance of fatty acid β-oxidation proteins, increased lactate levels and high abundance of glutamine around day 150-200, which could indicate a change in the functional properties of astrocytes at later time points. This could be a result of metabolic support provided by astrocytes to neurons. It could also indicate that this metabolic shift in astrocytes during development is needed to promote energy-demanding signaling in neurons such as synaptic release of neurotransmitters. We observed that levels of proteins related to oxidative phosphorylation increased at later time points, indicating that neurons gradually differentiated into more mature states in parallel to the appearance of differentiating astrocytic cells. Our study is however limited by the absence of oligodendrocytes in CBOs, which likely require longer culture durations to develop. The differentiation of oligodendrocyte precursors has been identified in a few previous studies [19, 92] and as oligodendrocytes develop *in vivo* at late fetal and postnatal stages [93], it could reflect the developmental stages currently represented in neural organoid models.

Comparative proteomic analysis revealed similarities between the proteomes of CBOs and human fetal brain tissue, underscoring the physiological relevance of organoids as robust models for studying human brain development. This concordance not only validates CBOs as a valuable tool for developmental neuroscience but also highlights their capability to recapitulate distinct periods associated with specific cortical structures and zones. Such insights are crucial for the strategic planning of experimental timelines and methodologies. Notably, progenitor cell functions specific to the ventricular zone are prominently modeled at the earliest time points in CBOs. As CBOs mature, the proteome shifts to reflect later developmental stages, where subplate and intermediate zone areas become more prominent. These zones are associated with critical processes in neuronal development, such as migration, axonal growth, and synapse development. The dynamic proteomic changes observed in CBOs parallels the sequential development of cortical zones in the human brain. By understanding these stages, researchers can align their experimental designs with specific neurodevelopmental milestones, thereby increasing the relevance of findings obtained from CBO models.

In conclusion, we performed a high temporal resolution protein-centric analysis of the abundance of proteins, PTMs and metabolites in CBOs covering periods of progenitor cell differentiation, neurogenesis, neuronal maturation and gliogenesis, which serve as a critical resource for the neural organoid field. Our findings demonstrate that temporal changes in protein and PTM abundances are related to neuronal differentiation and energy metabolism, whereas calcium signaling is mainly regulated on the post-translational level. This underlines the importance of PTMs in regulation of brain developmental events. We further show that the proteome of CBOs and human fetal brain is comparable, validating CBOs as a model for studying mechanisms of human brain development.

## Methods

### Pluripotent stem cell culture

The female IMR90 (clone #4) induced pluripotent stem cell line (iPSC) and the female H9 embryonic stem cell line (ESC) was obtained from WiCell. The cell lines were maintained in feeder-free conditions on Matrigel coated (GFR, Corning) cell culture dishes using mTESR1 medium (StemCell Technologies) at 37°C in 5% CO_2_ with daily medium change. Cells were passaged using Gentle Cell Dissociation Reagent (StemCell Technologies). H9 ESCs were approved for use in this project by the Scientific Ethics Committee from the Region of Southern Denmark (no. S-20210151).

### Organoid differentiation

Unguided neural organoids were generated based on the principles outlined by Lancaster et al. [1], and Lancaster and Knoblich [2] using the commercially available STEMdiff™ Cerebral Organoid Kit from StemCell Technologies. Organoids were generated according to the guidelines from the manufacturer with a few modifications enabling large-scale culturing of appropriate organoid numbers (Figure S1). On day 0, PSCs were dissociated to single cells using Gentle Cell Dissociation Reagent (StemCell Technologies) and plated in AggreWell™800 culture plates (StemCell Technologies) with 600,000 cells per AggreWell in EB formation medium containing Y-27632 ROCK inhibitor (50 mM, StemCell Technologies). From day 2-5, EBs were kept in EB Formation Medium. From day 5-7, EBs were kept in Neural Induction Medium. On day 7, early organoids were transferred to cell culture dishes and embedded in Matrigel (Corning). Organoids were embedded in batches of approximately 30 organoids by suspending and plating the organoids in a 60% ice-cold Matrigel (Corning) / 40% Expansion Medium mixture on cell culture dishes (around 10 mm diameter) using wide-bore pipette tips. Matrigel (Corning) was polymerized for 30 min at 37°C before adding Expansion Medium to the dishes. On day 10, the medium was exchanged to Maturation Medium containing 1% Penicillin-Streptomycin (Gibco) and 1 mg/ml Amphotericin B (Thermo Scientific) with biweekly medium changes until day 35. From day 10 onwards, the organoids were cultured on an orbital shaker (INFORS HT Celltron orbital shaker, 57 rpm). On day 15, the organoids were released from the Matrigel embedding by gently pipetting with wide-bore pipette tips. At day 35, the medium was exchanged to Improved Differentiation Medium + Vitamin A (IDM+A), containing a 1:1 mixture of DMEM/F12 (Gibco) and Neurobasal medium (Gibco) supplemented with 1% Glutamax (Gibco), 1% B27 + vitamin A (Gibco), 0.5% N2 supplement (Gibco), 0.5% MEM-NEAA (Gibco), 0.025% Insulin (Sigma), 50 mM 2-mercaptoethanol, 1% Penicillin-Streptomycin (Sigma) and 1 mg/ml Amphotericin B (Thermo Scientific). From day 35-70, 1% dissolved Matrigel (Corning) was added to the medium. Organoids were sliced into two equal parts by cutting them in the middle at day 43 to prevent hypoxia.

Each cell line was differentiated and organoids collected for profiling at the following 28 time points: day 17, D19, D21, D23, D25, D27, D30, D35, D40, D45, D50, D55, D60, D65, D70, D80, D90, D100, D110, D120, D130, D140, D150, D160, D170, D180, D190, and D200. Each cell line was subjected to three independent differentiations, adding up to a total of 84 samples per cell line. At the time of collection, organoids were washed in 50 mM ammonium acetate (pH 7.5) and subsequently kept at -70°C until sample processing.

### Sample processing for metabolomics, proteomics and PTMomics

#### Protein and metabolite extraction

Organoids were lysed in a lysis buffer (pH 7.5) consisting of 10 mM ammonium acetate, 3% SDC and PhosSTOP™ (Roche) by sonication at 40% amplitude for 4x10 sec. Samples were heated for 5 min at 110°C and subsequently centrifuged at 20,000 xg for 20 min at room temperature (RT) to pellet insoluble material. From the supernatant, 75 mg (samples from day 17-25) or 150 mg (samples from day 27-200) of protein was transferred to an Amicon Ultra-0.5 Centrifugal filter with a 10 kDa cutoff (Merck Millipore) and each sample was diluted to 1% SDC using a 10 mM ammonium acetate solution (pH 7.5). The solution was passed through the filter and metabolites present in the flow-through were transferred to a Captiva non-dripping 0.45 μm filter plate (Agilent Technologies). SDC was precipitated with 2% formic acid (FA) and the flow-through was centrifuged from the filter plate into low-binding 96-well plates to collect samples for metabolomics analysis. The metabolite samples were dried by vacuum centrifugation before LC-MS analysis.

#### CysPAT labelling of free cysteine residues

Cysteine-specific Phosphonate Adaptable Tag (CysPAT) was synthesized as described by Huang et al. [32] using 2 mg N-succinimidyl Iodoacetate (SIA) per 10 ml dimethyl sulfoxide (DMSO) and 2 mg 2-aminoethylphosphonic acid (2-AEP) per 160 ml 50 mM TEAB solution. The reaction between SIA and 2-AEP to generate CysPAT (2-(2-iodoacetamido)ethyl) took place in the dark under rotation for 2h. Proteins retained on top of the 10 kDa cutoff Amicon Ultra Centrifugal filters were dissolved in 100 mM HEPES buffer (pH 8.5) containing 1% SDC and the CysPAT reagent was added to each sample at a 5 mM concentration and left to react with the free cysteine residues in the proteins for 1 h at 37°C. After incubation, the solution was passed through the filter, and the proteins left on the filter were washed twice using a 100 mM HEPES buffer containing 0.2% SDC (pH 8.5).

#### Reduction and protein digestion

After the last wash, the proteins on the filter were dissolved in 100 mM HEPES buffer containing 1% SDC and 3 mM tris(2-carboxyethyl)phosphine (TCEP; pH 8.5) and incubated for 30 min at RT to reduce disulfide bonds between cysteine residues. Proteins were digested using lysyl endoproteinase (0.00004 AU per mg protein; Wako) and 5% (w/w) dimethylated trypsin [94] overnight (ON) at 37°C. After incubation, another 1% (w/w) dimethylated trypsin was added per sample and incubated for 1 h at 37°C. The digested peptides were transferred to a low-binding tube, and the filter washed with 30% acetonitrile and pooled with the peptide digest before each sample was dried by vacuum centrifugation.

#### TMTpro 16-plex labelling

The peptide samples were resolubilized in 100 mM HEPES buffer (pH 8.5) and equal amounts of each sample were used to generate a pooled sample containing peptides from all conditions. To ensure accurate quantitation between samples collected at different time points of organoid maturation, we labelled peptide samples using isobaric tandem mass tags according to the manufacturer’s instructions. A total of 12 sets of a TMTpro 16-plex (Thermo Fisher Scientific) were used to label the 168 samples. In each TMT set, one channel was used for labelling of the pooled sample to enable comparison of samples between the different TMT sets. The labelling reaction was checked by LC-MS/MS analysis to ensure proper labelling of all TMT channels in all TMT sets. Subsequently, samples were pooled in 12 separate TMT sets in a 1:1 ratio, and excess reagent was quenched using 10 ml of a 1 M ammonia bicarbonate buffer by incubation for 15 min at RT. After incubation, the pooled TMT samples were acidified with 2% FA and vortexed to pellet SDC. The samples were centrifuged at 20,000 xg for 15 min at RT and the supernatant was transferred to a new low-binding tube and dried by vacuum centrifugation to a volume of 150 ml.

### Enrichment of various cysteine-containing peptides and PTM peptides

#### Enrichment of phosphorylated peptides, sialylated N-glycopeptides and CysPAT labeled free cysteine-containing peptides (SIA)

For all 12 TMT samples the partially dried peptide solution was diluted to 1 ml 5% TFA, 1 M glycolic acid and 80% acetonitrile. To each sample, a total of 10 mg of TiO_2_ beads (GL Science, Japan) was added and the solution incubated at RT with gentle rotation for 10 min. After incubation, the sample was centrifuged for 15 sec to pellet the TiO_2_ beads. The supernatant was transferred to another Eppendorf tube and 5 mg TiO_2_ beads was added and incubated with rotation at RT for 10 min. After incubation, the sample was centrifuged in a table centrifuge and the supernatant was recovered in a new Eppendorf tube. The beads from both incubations were pooled with 200 µl 1% trifluoroacetic acid (TFA), 80% acetonitrile in a new Eppendorf tube and after mixing and centrifugation, the supernatant was added to the first supernatant. This supernatant was lyophilized by vacuum centrifugation. The washed TiO_2_ pellet was dried by vacuum centrifugation for 10 min and the beads were resolubilized in 150 µl 100 mM HEPES, pH 8.5. PNGaseF and Sialidase A were added to the solution, and the solution was mixed and incubated at 37°C ON. After incubation, the sample was diluted to 1% TFA and 70% acetonitrile and mixed with rotation for 10 min to allow the phosphopeptides and SIA peptides to reattach to the TiO_2_ beads. After incubation, the sample was centrifuged at 14,000g to pellet the TiO_2_ beads and the supernatant was transferred to an Eppendorf tube named “Deglycopeptides” and lyophilized. A total of 150 µl 5% ammonia water was added to the TiO_2_ pellet and the sample was mixed well and left at RT for 10 min to allow efficient elution of phosphopeptides and SIA peptides. After incubation, the TiO_2_ beads were pelleted by centrifugation and the supernatant was transferred to a p200 tip with a C8 disc and the solution was pushed through the C8 disc to capture TiO_2_ beads. The solution was recovered in a low-binding Eppendorf tube. A total of 50 µl 5% ammonia water and 50 µl acetonitrile was added to the TiO_2_ beads and the sample was mixed and centrifuged. The supernatant was pushed through the C8 disc and collected into the other elution. The final eluted phosphopeptides and SIA peptides were lyophilized prior to high pH reversed-phase (RP) fractionation.

The first supernatant from the TiO_2_ enrichment was resolubilized in 2 ml 0.1% TFA and peptides were desalted using a 5 mg HLB cartridge (Waters, UK). After washing, the peptides were eluted with 1 ml 70% acetonitrile in water and the solution was lyophilized.

#### Purification of cysteine-containing peptides with reversible modifications

After lyophilization, the peptide solution was resolubilized in 50 µl 500 mM HEPES, pH 8.5 and TCEP was added to 5 mM and the solution was incubated for 1 h at RT. During incubation, a total of 300 µl Agarose, S3 high-capacity acyl-rac capture beads (NANOCS Inc, US) were washed twice in 100 mM HEPES, pH 8.5. After TCEP reduction, the peptide solution was rapidly diluted to 1 mM in H_2_O and the S3 beads were added. The solution was incubated at RT for 1 h with rotation. After incubation, the beads were pelleted by gentle centrifugation using a table centrifuge and the supernatant was transferred to another Eppendorf tube labelled “S3 FT”. The S3 beads were washed with 300 µl 100 mM HEPES, pH 8.5 and the supernatant was added to the S3 FT fraction. The S3 beads were solubilized in 500 µl H_2_O and transferred to a MobiSpin column filter (10 μm, MoBiTec Molecular Biology, Germany) and the beads were washed 3 times with 500 µl H_2_O. The S3 beads were mixed with 300 µl 100 mM HEPES, pH 8.5 containing 10 mM DTT to release the cysteine-containing peptides and incubated at RT for 30 min. After incubation, the solution was spun through the MobiSpin column filter and the reversibly modified cysteine-containing peptides (RmCys peptides) were collected in a new Eppendorf tube, alkylated with 20 mM iodoacetamide for 30 min at RT in the dark and finally lyophilized prior to high pH RP fractionation.

#### Enrichment of lysine-acetylated peptides

The S3 FT fraction was adjusted to pH 7.2 using hydrochloric acid (HCl) and a total of 20 µl Immunoaffinity beads carrying Anti-Lysine-acetyl antibodies conjugated to Agarose beads (PTM-Scan^®^ Acetyl-Lysine Motif [Ac-K], Cell Signalling Technology), and the solution was incubated for 2 h at RT with gentle rotation. After incubation, the beads were pelleted by gentle centrifugation and the supernatant was transferred to a new Eppendorf tube labelled “non-modified peptides”. The beads were washed using the buffer in the PTMScan kit three times and finally, the lysine-acetylated peptides were eluted with 150 µl 0.15 % TFA for 10 min. After incubation, the beads were pelleted, and the supernatant transferred to another Eppendorf tube, and the beads were washed with 100 µl 0.15 % TFA. The supernatant from the wash was mixed with the first supernatant and lyophilized prior to LC-MS/MS.

#### High pH Reversed-Phase (RP) fractionation

The high pH RP separation was performed essentially as described in Ashrafian et al. [95]. The phosphopeptides and SIA peptides were separated into 20 concatenated fractions. The RmCys peptides were fractionated into 12 concatenated fractions. The non-modified peptides were separated into 20 concatenated fractions. All the fractions were lyophilized prior to LC-MS/MS.

### LC-MS analysis of metabolites

Metabolite samples were resuspended in 25 µl 0.1% FA of which 3 µl were transferred to a common quality control (QC) sample. A total of 4 µl of each sample was injected using a Vanquish Horizon UPLC (Thermo Fisher Scientific, Germering, Germany) equipped with a Zorbax Eclipse Plus C18 guard (2.1 × 50 mm and 1.8 µm particle size, Agilent Technologies, Santa Clara, CA, USA) and an analytical column (2.1 × 150 mm and 1.8 µm particle size, Agilent Technologies, Santa Clara, CA, USA) kept at 40°C. The flow rate was set to 400 µl/min using eluent A (0.1% FA) and eluent B (0.1% FA, acetonitrile): 3% B from 0 to 1.5 min, 3-40% B from 1.5 to 4.5 min, 40☐95% B from 4.5 to 7.5 min, 95% B from 7.5 to 10.1 min and 95 to 3% B from 10.1 to 10.5 min before equilibration for 3.5 min with the initial conditions. The flow was coupled to a Q Exactive HF mass spectrometer (Thermo Fisher Scientific, Bremen, Germany) for mass spectrometric analysis in both positive and negative ion mode using the following general settings for MS1 mode: resolution: 120,000, AGC target: 3e6, maximum injection time: 200 ms, scan range 65-975 m/z and lock mass: 391.28429/112.98563 (pos/neg mode). For compound fragmentation MS1/ddMS2 mode was used with the following general settings: resolution: 60,000/15,000, AGC target: 1e6/1e5, maximum injection time: 50/100 ms, scan range 65-975 m/z, loop count: 10, isolation width: 2 m/z and normalized collision energy: 35/38 (pos/neg).

### LC-MS/MS analysis of non-modified peptides

The non-modified peptides from the high pH RP separations were all analyzed by SPS-RT-MS^3^ [96]. The 20 concatenated fractions from each high pH RP separation were resolubilized in 4 µl 0.1% FA and 3.5 µl was loaded onto a 20 cm 75 μm inner diameter (ID) column containing C18 material (Reprosil 1.9 µm, Dr. Maisch, Ammerbuch-Entringen, Germany) using a nano-Easy LC (Thermo Fisher Scientific) coupled to an Orbitrap Eclipse Tribrid mass spectrometer (Thermo Fisher Scientific). The peptides were eluted using a 120 min gradient (2-25% buffer B (90% ACN in 0.1% FA) over 100 min, 25-40% buffer B over 20 min and finally 40-95% buffer B in 1 min). MS1 spectra were recorded for m/z range 350-1600 using 120,000 FWHM resolution using an AGC value of 250% and maximum filling time of 50 ms. Peptides were selected for 3 seconds cycle time for MS/MS using an isolation window of 0.7 Da, and peptides were fragmented using CID (NCE 35) in the linear ion trap using turboscans. Each CID low resolution spectra were searched using real time searching in the building database software using the following settings: Human Uniprot fasta database (version from 2023.06.28); TMTpro (K and N-terminal) as fixed modifications; mass accuracy 10 ppm for MS1 and 20 ppm for MS2. For the precursors who matched a sequence in the MS2 search, the precursor was reselected and fragmented and 10 fragment ions were selected by synchronous precursor selection (SPS) for HCD fragmentation (NCE 55) and the TMT reporter ions were recorded in the orbitrap with TurboTMT settings.

### LC-MS/MS analysis of PTM- and cysteine-containing peptide samples

#### Phosphorylated/SIA peptides and RmCys peptides

The peptide fractions from each high pH RP separations for the phosphorylated/SIA peptide and RmCys peptide analyses were resolubilized in 4 µl 0.1% FA and 3.5 µl was loaded onto a 20 cm 75 μm ID column containing C18 material (Reprosil 1.9 µm, Dr. Maisch, Ammerbuch-Entringen, Germany) using a nano-Easy LC (Thermo Fisher Scientific) coupled to an Orbitrap Exploris 480 mass spectrometer (Thermo Fisher Scientific). The phosphorylated/SIA peptides (20 concatenated fractions per TMT set) were eluted using a 90 min gradient (2-25% buffer B (90% ACN in 0.1% FA) over 75 min, 25-40% buffer B over 15 min and finally 40-95% buffer B in 1 min). The RmCys peptides (12 concatenated fractions per TMT set) were eluted using a 120 min gradient (2-25% buffer B (90% ACN in 0.1% FA) over 100 min, 25-40% buffer B over 20 min and finally 40-95% buffer B in 1 min). MS1 spectra were recorded for m/z range 350-1400 using 120,000 FWHM resolution using an AGC value of 300% and maximum filling time of 50 ms. Peptides were selected for 3 seconds cycle time for MS/MS using an isolation window of 0.7 Da, HCD with NCE 33, a resolution of 45,000 FWHM, AGC target value of 300% and maximum injection time of 86 ms.

#### Deglycopeptides and lysine-acetylated peptides

For the deglycosylated formerly sialylated N-glycopeptides and the lysine-acetylated peptides, the samples were dissolved in buffer A (0.1% FA) and analyzed by nano LC-ESI-MS/MS using an EASY-nLC (Thermo Fisher Scientific) connected to an Orbitrap Fusion Lumos mass spectrometer (Thermo Fisher Scientific, USA). The separation was performed on an in-house-made fused silica capillary two-column setup, a 3 cm pre-column (100 μm inner diameter packed with Reprosil-Pur 120 C18-AQ, 5 μm (Dr. Maisch GmbH)), and an 18 cm pulled emitter analytical column (75 μm ID packed with Reprosil-Pur 120 C18-AQ, 3 μm (Dr. Maisch GmbH)). The peptides were eluted using a 120 min gradient (2-25% buffer B (90% ACN in 0.1% FA) over 100 min, 25-40% buffer B over 20 min and finally 40-95% buffer B in 1 min). MS1 spectra were recorded for m/z range 350-1200 using 120,000 FWHM resolution using an AGC value of 250% and maximum filling time of 50 ms. Peptides were selected for 3 seconds cycle time for MS/MS using isolation window of 0.7 Da, HCD with NCE 34, a resolution of 50,000 FWHM, AGC target value of 300% and maximum injection time of 86 ms. The deglycosylated formerly sialylated N-glycopeptides were analyzed in technical duplicates, whereas the lysine-acetylated peptides were analyzed in one injection.

### Data processing

#### Metabolomics

Raw data was processed with MzMine (v 3.9) [97]. In brief, the following modules were used: Mass detection, ADAP chromatogram builder, ADAP chromatogram resolver, 13C Isotope filter, Join aligner, and Gap filling (same RT and m/z range). Compounds were annotated at Metabolomics Standards Initiative (MSI) level 3 [98] using local MS/MS spectra databases of National Institute of Standards and Technology 17 (NIST17), Mass Bank of North America (MoNA) and all spectral libraries available from MS-DIAL (v Aug., 2022) [99]. After compound annotation, the datasets were filtered and normalized in MetaboLink (https://github.com/anitamnd/MetaboLink/wiki-manuscript submitted to Bioinformatics). Finally, the signals were auto-scaled and log2-transformed in MetaboAnalyst [100].

#### Proteomics

Raw data files were searched in Proteome Discoverer (version 2.5.0.400, Thermo Scientific) using the Sequest HT search engine [101]. The raw files were searched against the human proteome database (version from 2023.06.28). During the database search, the following parameters were used: trypsin was set as enzyme and allowed for maximum 2 missed cleavages with TMTpro on the peptide N-terminus and lysine residues included as fixed modifications. Precursor mass tolerance was set to 10 ppm and fragment mass tolerance was set to 0.8 Da. Peptide identifications were filtered against a 1% false discovery rate cut-off using the Percolator algorithm [102]. Quantification was performed based on TMT reporter ion intensities during MS3 scans. Peptide quantifications were rolled up to protein groups and quantification across different sets of TMTpro 16-plex was normalized based on a common reference channel containing a mix of all samples. Only proteins with 2 or more unique peptides identified were considered for further analysis.

#### PTMomics

Raw data files from each of the PTM-enriched peptide data sets were searched in Proteome Discoverer (version 2.5.0.400, Thermo Scientific) using the Sequest HT search engine [101]. The raw files were searched against the human proteome database (version from 2023.06.28). During the database searches, the following parameters were used: trypsin was set as enzyme and allowed for maximum 2 missed cleavages (up to 4 missed cleavages were allowed for the lysine-acetylated peptides) and TMTpro on the peptide N-terminus as well as lysine residues were included as fixed modifications (TMTpro on lysine residues was considered a dynamic modification for the lysine-acetylated peptides). Dynamic modifications considered during the database search were acetylation of lysine residues for the lysine-acetylated peptides, carbamidomethylation of cysteine residues for the reversibly modified cysteine-containing peptides, SIA on cysteine residues and phosphorylation of serine, threonine, and tyrosine residues for the free cysteine- and phosphorylation-containing peptides, and deamidation of asparagine residues for the formerly sialylated glycopeptides. Precursor mass tolerance was set to 10 ppm and fragment mass tolerance was set to 0.05 Da. Peptide identifications were filtered against a 1% false discovery rate cut-off using the Percolator algorithm [102]. Quantification was performed based on TMT reporter ion intensities during MS2 scans. Quantification across different sets of TMTpro 16-plex was normalized based on a common reference channel containing a mix of all samples. For each of the datasets, peptides that did not carry the modification of interest were excluded from the subsequent analysis. Thus, lysine-acetylated peptides had to carry acetylation of a lysine residue, reversibly modified cysteine-containing peptides had to have a carbamidomethylated cysteine residue, free cysteine-containing peptides had to carry SIA from the CysPAT tag, phosphorylated peptides had to carry phosphorylation on a serine, threonine, or tyrosine residue, and formerly sialylated N-glycopeptides had to have a deamidated asparagine residue in the peptide sequence. As spontaneous deamidation can occur during the sample preparation process, we further filtered the formerly sialylated N-glycopeptides for the N-linked glycosylation motif (N-X-S/T/C, X≠P), and only accepted peptides known to be N-glycosylated according to the UniProt Knowledgebase.

### Data analysis

Each dataset was subjected to a filtering step to include features identified in at least 20% of the samples in the subsequent analyses. Normalized abundance values were log-transformed, and missing values were imputed using a k-nearest neighbor algorithm with k=5 neighbors implemented in the VIM R package [103]. To enable comparison across datasets, abundance values were z-transformed. Principal component analysis was performed using the prcomp() function from the stats package in R and the first two principal components were visualized using the factoextra R package [104]. Correlation analysis was performed between the first 5 principal components and study covariates such as time point, cell line, and batch number using the corrplot R package [105]. Statistical comparisons were performed for all time points against the first time point (D17) to identify features with altered abundance during organoid maturation. For the proteomics and modified peptide datasets, a PolySTest [106] was performed, whereas for the metabolomics data, a Welch’s two-sample t-test with Benjamini-Hochberg correction [107, 108] was applied. Features with an absolute log_2_ fold change from D17 of minimum 1 and p_adj_<0.05 in at least one time point were considered significantly altered during the time course.

#### Cluster analysis

Features with altered abundance during organoid maturation were subjected to unsupervised k-means clustering. For this, the protein-related datasets were divided into two categories: reflectors of protein abundance (proteomics, reversibly modified cysteine-containing peptides, and formerly sialylated N-glycopeptides) and reflectors of dynamic protein modifications (phosphorylation, free cysteine-containing peptides, and lysine acetylation). Optimal k-means clusters for each data category were determined based on the elbow method, silhouette coefficient, and gap statistics using the factoextra R package [104] (figure S5). For protein abundance, the optimal number of clusters was 10, and for dynamic protein modifications, the optimal number of clusters was 11. For protein abundance and dynamic protein modification clusters, the protein members of each cluster were subjected to KEGG pathway and Gene Ontology cellular component enrichment using the compareCluster() function from the clusterProfiler package in R [109, 110] with Benjamini-Hochberg correction for multiple testing. All proteins identified across the proteomics and modified peptide datasets were used as background for the enrichment analysis. Enriched KEGG terms related to diseases and organ systems other than the central nervous system were excluded. To avoid visualization of redundant KEGG pathway terms, one KEGG term from each KEGG pathway subcategory was selected. The selection was based on q-value after enrichment (lowest and/or lower than 0.01) and the highest count. Top 5 Gene Ontology Cellular Component terms per cluster were visualized. From the enriched KEGG pathway terms ‘axon guidance’, ‘calcium signaling pathway’, ‘motor proteins’, and ‘fatty acid degradation’, mapped proteins in clusters where the term was found to be enriched were visualized in protein-protein interaction diagrams using the STRING database [111]. Protein-protein interaction diagrams were drawn in Cytoscape using the stringApp (v2.0.1) [112] and Omics Visualizer (v1.3.0) [113] apps. Protein-protein interactions with high confidence (>0.7) were visualized.

#### Comparison to human brain tissue proteome

The proteome of CBOs from day 17-200 identified by proteomics analysis was compared to FFPE human brain tissue proteomes from the ventricular zone, intermediate zone, subplate, and cortex regions analyzed by Djuric et al. [68]. We only considered proteins with more than 2 unique peptides identified and proteins without missing values between different human brain samples for correlation analysis with CBO samples. This left 1,134 proteins from the human brain samples, of which 1,025 proteins were also identified in CBOs and used for correlation analysis. Pearson correlation coefficients between human brain and CBO samples were calculated and visualized using the corrplot package in R [105].

### Multi-electrode array (MEA) analysis

Spontaneous electrical activity in organoids was examined using a BioCam Duplex MEA system (3Brain). Before attachment of the organoids to the recording area, the Accura HD-MEA cartridges (3Brain) were coated with 100 µg/ml poly-L-lysine (Sigma) ON. The cartridges were then washed 3 times with dPBS (Thermo Fisher) and coated with 50 µg/ml laminin (Sigma) ON. The following day, laminin was removed from the cartridge, and an organoid (day 104-106) was placed on the recording area using a sterile spatula followed by removal of the medium present around the organoid. To allow the organoid to attach to the recording area, small amounts of medium (IDM+A; 10-100µl/time) were slowly added every 10 minutes until the organoid had attached. Then, the cartridge was filled with a total of 1.5 ml medium. The organoids were cultured in the MEA cartridges with biweekly medium changes. Recordings of the spontaneous electrical activity of 10 individual organoids (from both cell lines and 4 independent differentiations) were performed on days 120, 140, 160, 180, and 200 on the BioCam Duplex MEA system. The recordings were each acquired for 2 min with spike detection in balanced mode and temperature set at 37°C. Analysis of the recordings was performed using the BrainWave 5 software (3Brain). For each organoid, electrode units were manually examined and units with neuronal spike activity were selected using the unit selection tool and grouped into a unit group. The units within each unit group were tracked across all time points. The mean firing rates across the different time points were compared using a repeated measures one-way ANOVA with Tukey’s multiple comparisons test.

### Immunohistochemistry

Organoids were fixed in 4% paraformaldehyde (Thermo Scientific) in PBS (Gibco) for 1 hour at RT. Fixed organoids were soaked in 30% sucrose in PBS at 4°C ON. Afterwards, organoids were embedded in an OCT-mounting medium and frozen in ice-cold ethanol below -50°C and subsequently stored at -70°C. CBOs were sectioned in 30 µm sections on a Leica CM1680 cryostat at -17 to -20°C. Sections were plated on Superfrost^®^ Plus glass slides and stored at -20°C.

The organoid slices were hydrated with PBS, washed 2x 5 min in 0.1% Saponin (Sigma) in PBS and incubated for 30 min with permeabilization and blocking buffer (5% goat serum and 0.1% Saponin in PBS). The slices were incubated ON at 4°C with primary antibodies in 5% serum and 0.1% Saponin in PBS: Mouse anti-microtubule-associated protein 2 (MAP2)ab 1:2000 (Sigma, M1406); rabbit anti-glial fibrillary acidic protein (GFAP) 1:3000 (Dako, Z0334); rabbit anti-Synapsin 1 (Syn1) 1:500 (Abcam, ab254349); Mouse anti-β-III-Tubulin 1:2000 (Sigma, T8660). The samples were washed 3x 10 min with 0.1% Saponin in PBS and incubated with a combination of secondary antibodies for 1h in the dark at RT: Goat anti-rabbit Alexa Fluor 647 1:1000, goat anti-rabbit Alexa Fluor 488 1:500, goat anti-mouse Alexa Fluor 568 1:1000 (all Invitrogen). The slices were washed 3 x 10 min with PBS and mounted with DAPI-containing ProLong Diamond Antifade mountant (Invitrogen). Slices were kept at 4°C in the dark until image acquisition. Images were aquired using a Nikon AX confocal unit on a Ti-2 LFOV microscope with equal settings between the timepoints analysed. The images shown were edited using ImageJ version 2.14.0.

### Western blotting

CBOs were lysed in 3% SDC in 100 mM HEPES, pH 8.5, sonicated 2 x 10 sec on ice at 40% amplitude and boiled for 5 min at 110°C. Following sonication, samples were centrifuged at 20000 rcf for 10 min at RT and supernatant transferred to new Eppendorf tubes. Protein concentration was determined by Nanodrop (Implen). Proteins were separated using Bolt 4-12% Bis-Tris Plus Gels (Thermo Scientific) with samples loaded in NuPage LDS Sample buffer (Thermo Scientific) with NuPage Sample Reducing agent (Thermo Scientific) and PageRuler Plus pre-stained protein standard (Thermo Scientific) for size reference. Proteins were transferred to PVDF membranes using the Trans-Blot® Turbo™ Transfer System (BioRad). Membranes were blocked with 5% skimmed milk in TBST buffer for 1 h and incubated ON at 4°C in 5% skimmed milk/TBST with the following primary antibodies: anti-mouse Map2a+b (Sigma #M1406) 1:500, anti-mouse ꞵ-actin HRP-linked (Abcam #ab49900) 1:50000, anti-rabbit Synapsin 1 (Abcam #ab254349) 1:1000, and anti-rabbit Glial Fibrillary Acidic Protein (GFAP, Dako #Z0334) 1:1000. Following 3 x wash in TBST, the membranes were incubated for 1 h at RT in 5% skimmed milk TBST with the following secondary antibodies: anti-rabbit IgG, HRP-linked (Cell signalling #7074) 1:10000 or anti-mouse IgG, HRP-linked (Abcam #ab6728) 1:10000. Following 3 x wash in TBST, the membranes were visualised with Immobilon ECL Ultra Western HRP Substrate (Millipore) using an Amersham 680 Imager (GE Healthcare). Blots were quantified using BioRad Image Lab (version 6.1).

### Data visualization

Illustrations were prepared using BioRender. Individual graphs of marker proteins and metabolites were prepared in R Studio and the average z-scaled abundance was fitted to a LOESS (locally estimated scatterplot smoothing) regression model [114] from the ggplot2 R package [115]. Other graphs were prepared using GraphPad Prism version 10.2.2 (GraphPad Software).

## Acknowledgements

We thank Arkadiusz Nawrocki for technical assistance. The Villum Center for Bioanalytical Sciences at the University of Southern Denmark is acknowledged for access to state-of-the-art mass spectrometry instrumentation.

