## Supplemental Figures for "Temporal proteomic and PTMomic atlas of cerebral organoid development"

### Supplementary figures

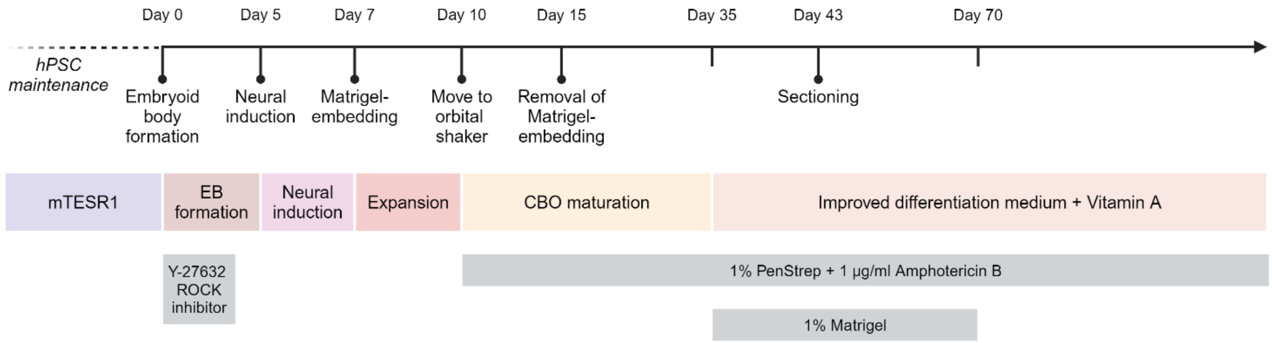


**Figure S1:** Schematic overview of the approach for culturing unguided forebrain organoids. The main steps during organoid formation, differentiation, and maturation and their timing are listed in the timeline. Medium type and duration are listed in colored boxes, whereas additional supplements are listed in grey boxes.

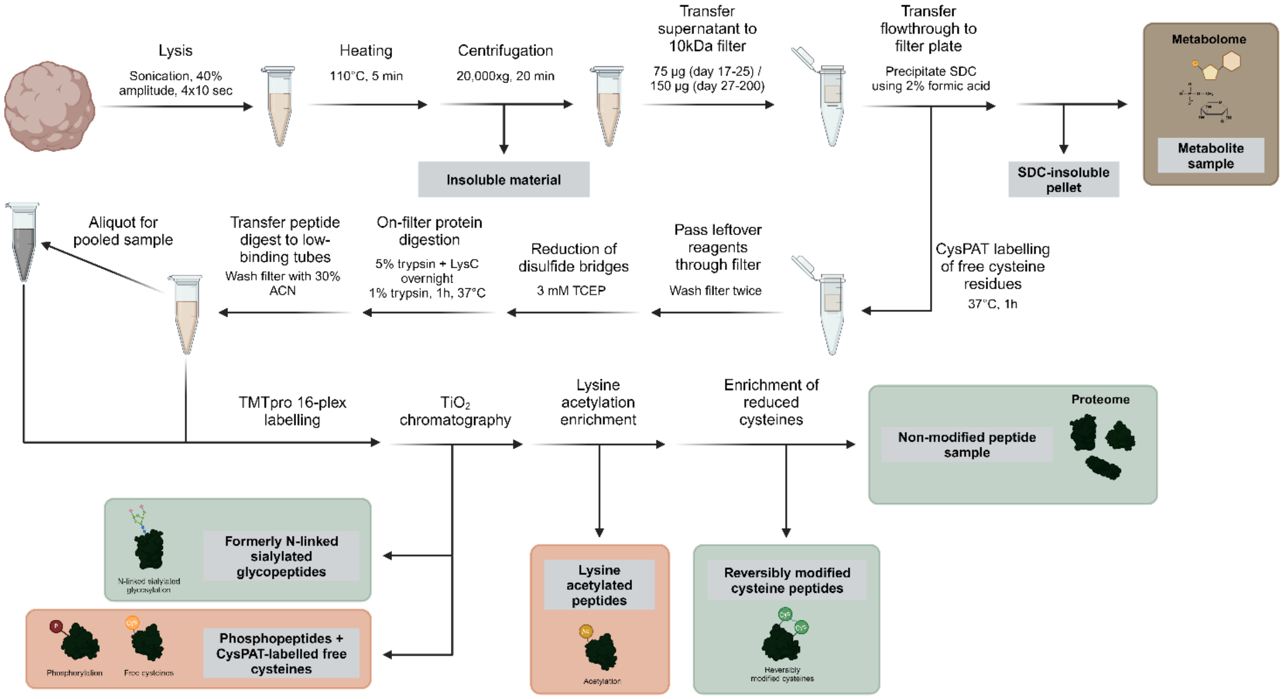


**Figure S2:** Schematic overview of the sample preparation workflow for multi-omic analysis of organoid samples. Samples collected for analysis are listed in colored boxes. Samples in green boxes reflect protein abundance (formerly N-linked sialylated glycopeptides, reversibly modified cysteine peptides, and non-modified peptides). Samples in orange boxes reflect dynamic protein modifications (phosphopeptides, CysPAT-labelled free cysteines, and lysine acetylated peptides). The metabolite sample is in a brown box.

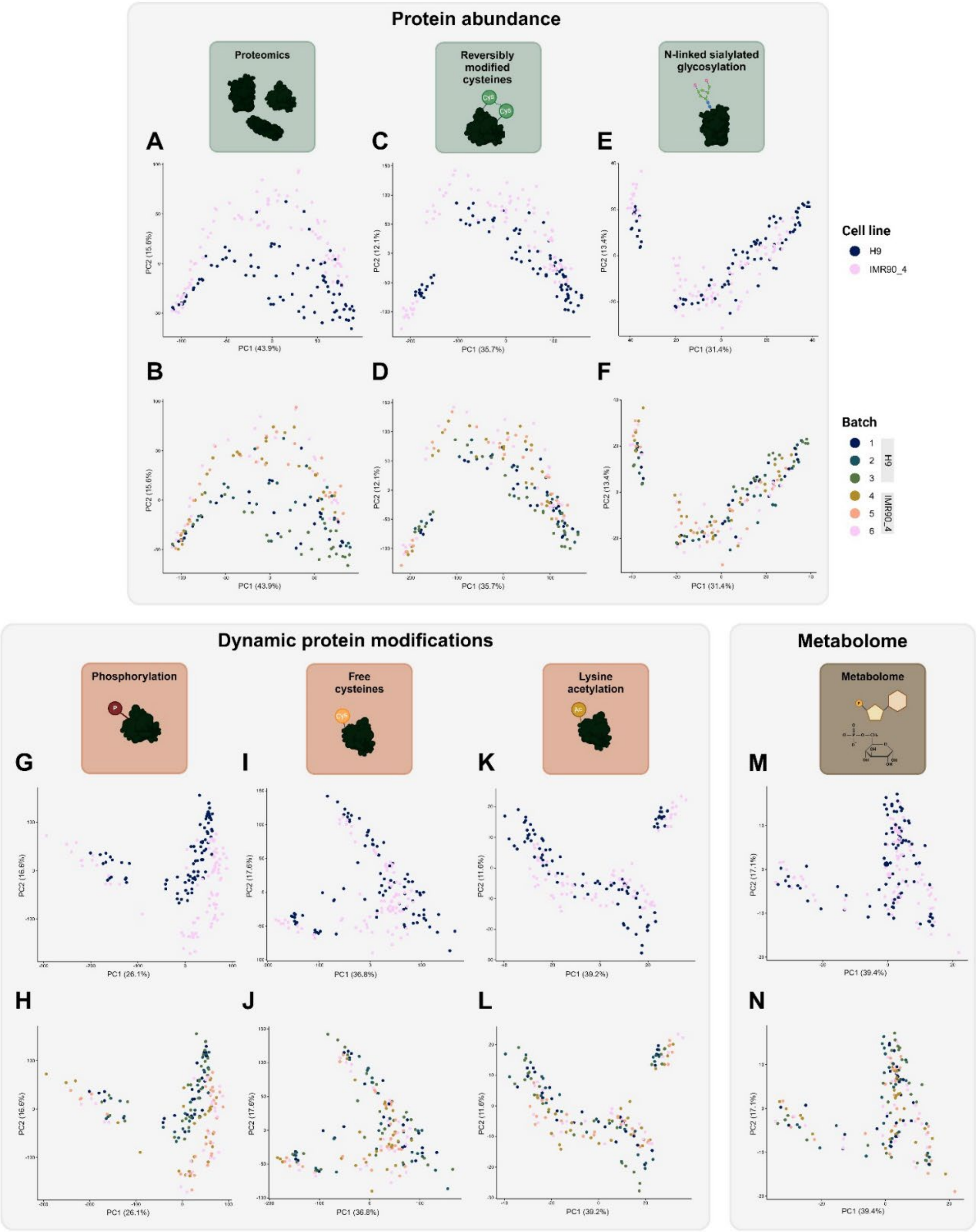


**Figure S3: A-F**: Principal component analysis of proteomics (**A+B**), reversibly modified cysteines (**C+D**), and N-linked sialylated glycosylation (**E+F**) data, which reflect temporal profiles of protein abundance. Samples are colored according to cell line in **A**, **C**, and **E**. Samples are colored according to individual organoid batch number in **B**, **D**, and **F**. **G-L**: Principal component analysis of phosphorylation (**G+H**), free cysteines (**I+J**), and lysine acetylation (**K+L**) data, which reflect temporal profiles of dynamic protein modifications. Samples are colored according to cell line in **G**, **I**, and **K**. Samples are colored according to individual organoid batch number in **H**, **J**, and **L**. **M-N**: Principal component analysis of metabolomics data. Samples are colored according to cell line in **M**, and according to individual organoid batch number in **N**.

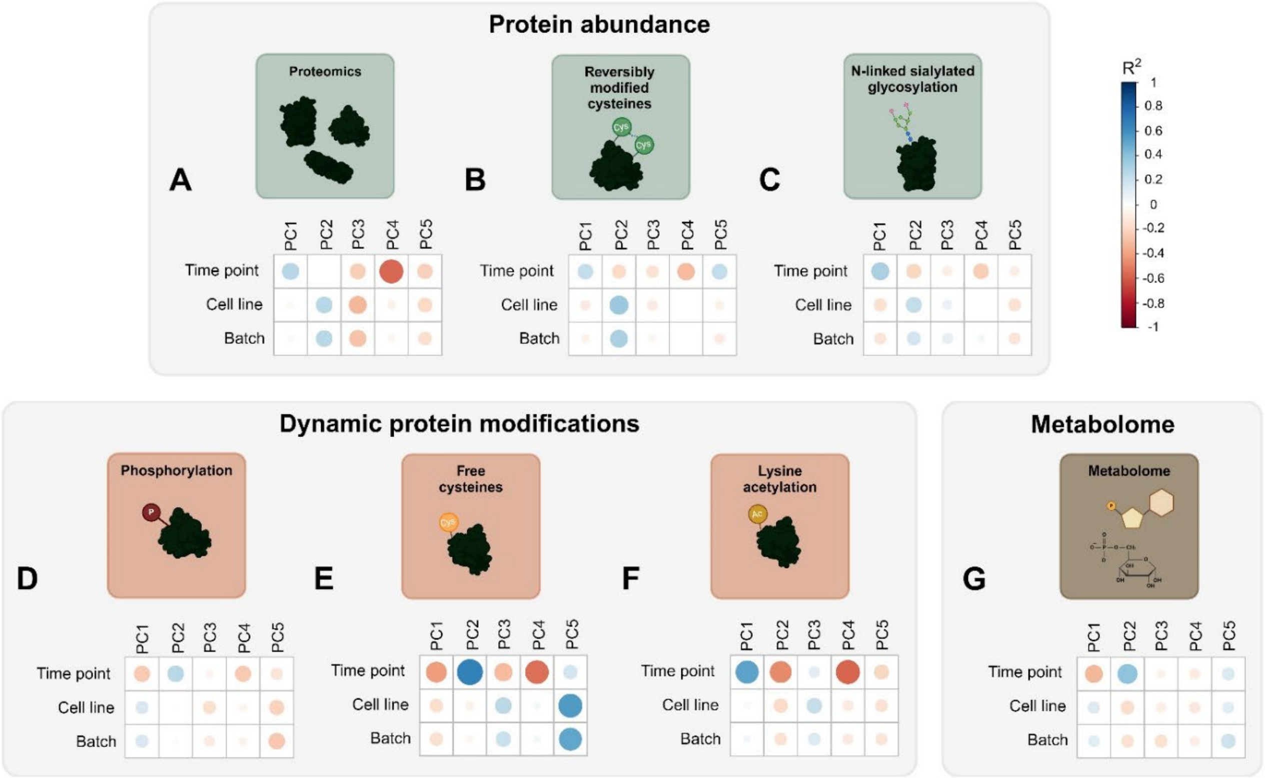


**Figure S4:** Correlation between covariates (time point, cell line, and batch) and the first 5 principal components from principal component analysis. **A-C:** Reflectors of protein abundance: proteomics (**A**), reversibly modified cysteines (**B**), and N-linked sialylated glycosylation (**C**). **D-F:** Reflectors of dynamic protein modifications: phosphorylation (**D**), free cysteines (**E**), and lysine acetylation (**F**). **G:** Metabolomics.

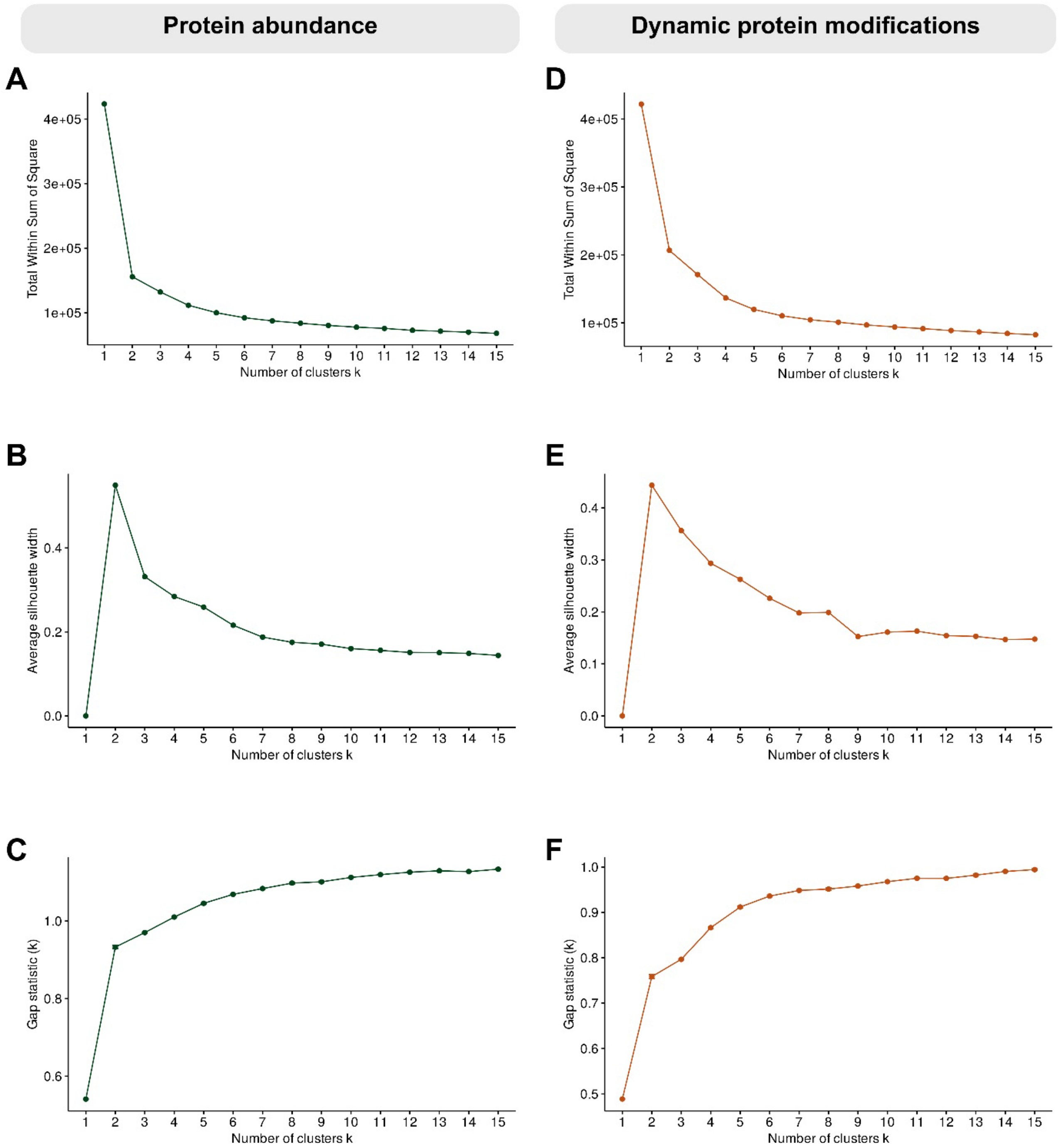


**Figure S5:** Results of estimation of optimal number of clusters for unsupervised k-means clustering of protein abundance and dynamic protein modification reflectors. **A:** Total within sum of square for protein abundance reflectors. **B:** Average silhouette width for protein abundance reflectors. **C:** Gap statistics for protein abundance reflectors. **D:** Total within sum of square for dynamic protein modification reflectors. **E:** Average silhouette width for dynamic protein modification reflectors. **F:** Gap statistics for dynamic protein modification reflectors.

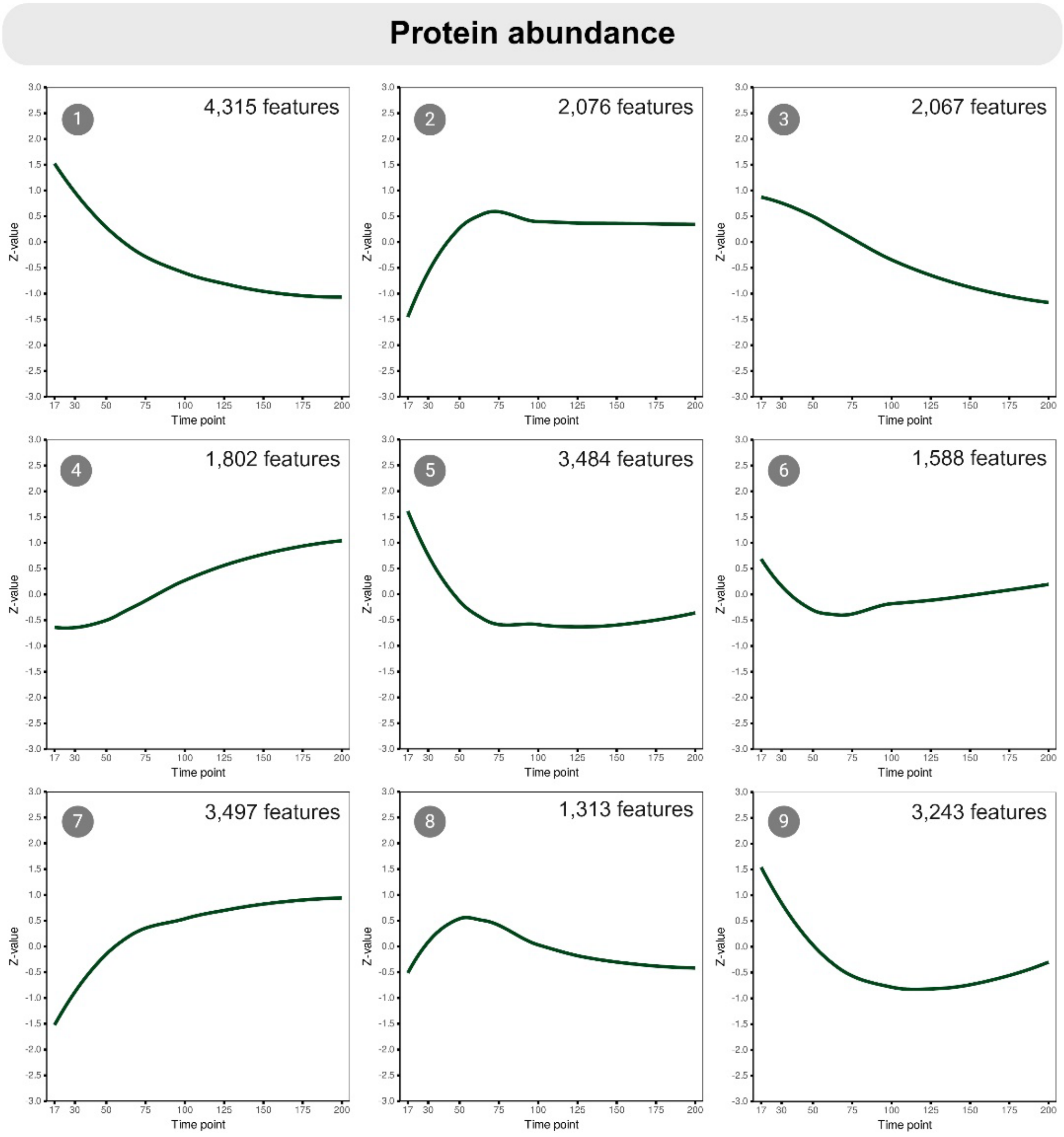


**Figure S6:** Plot of the average z-scaled abundance profile of each protein abundance cluster. Average z-scaled abundances were fitted to a LOESS (locally estimated scatterplot smoothing) regression model.

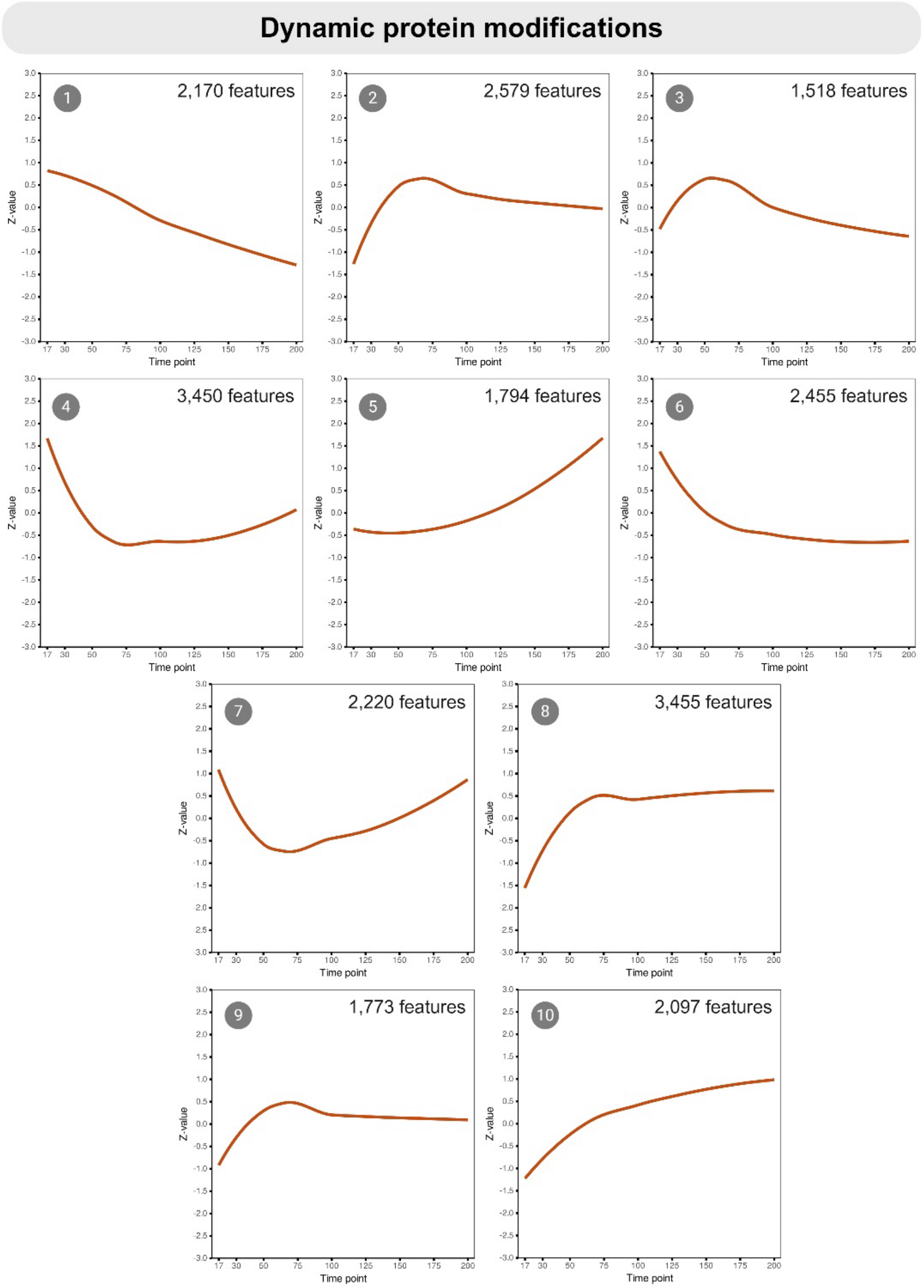


**Figure S7:** Plot of the average z-scaled abundance profile of each dynamic protein modification cluster. Average z-scaled abundances were fitted to a LOESS (locally estimated scatterplot smoothing) regression model.

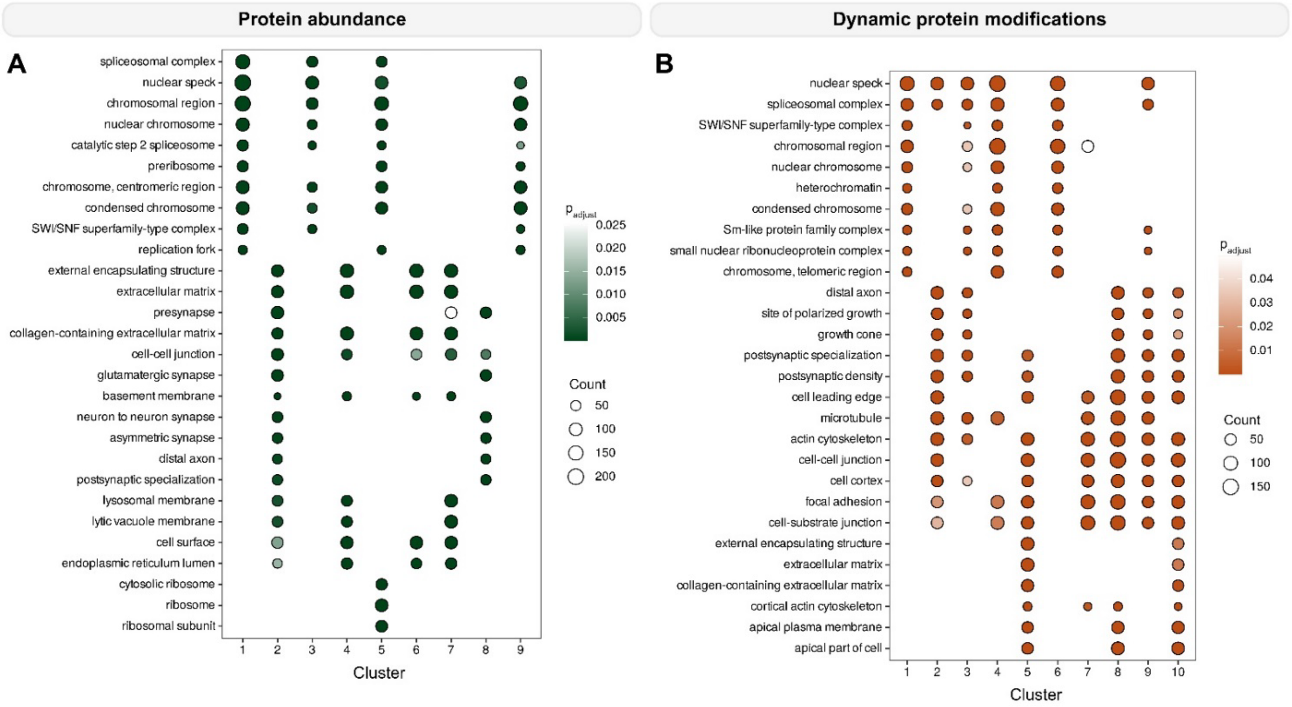


**Figure S8:** Enrichment results of Gene Ontology Cellular Components for protein abundance clusters (**A**) and dynamic protein modification clusters (**B**). The top 5 cellular components are shown for each cluster. Circle size reflects the number of proteins identified in a cluster, whereas circle color reflects enrichment p-value after Benjamini-Hochberg adjustment for multiple testing.

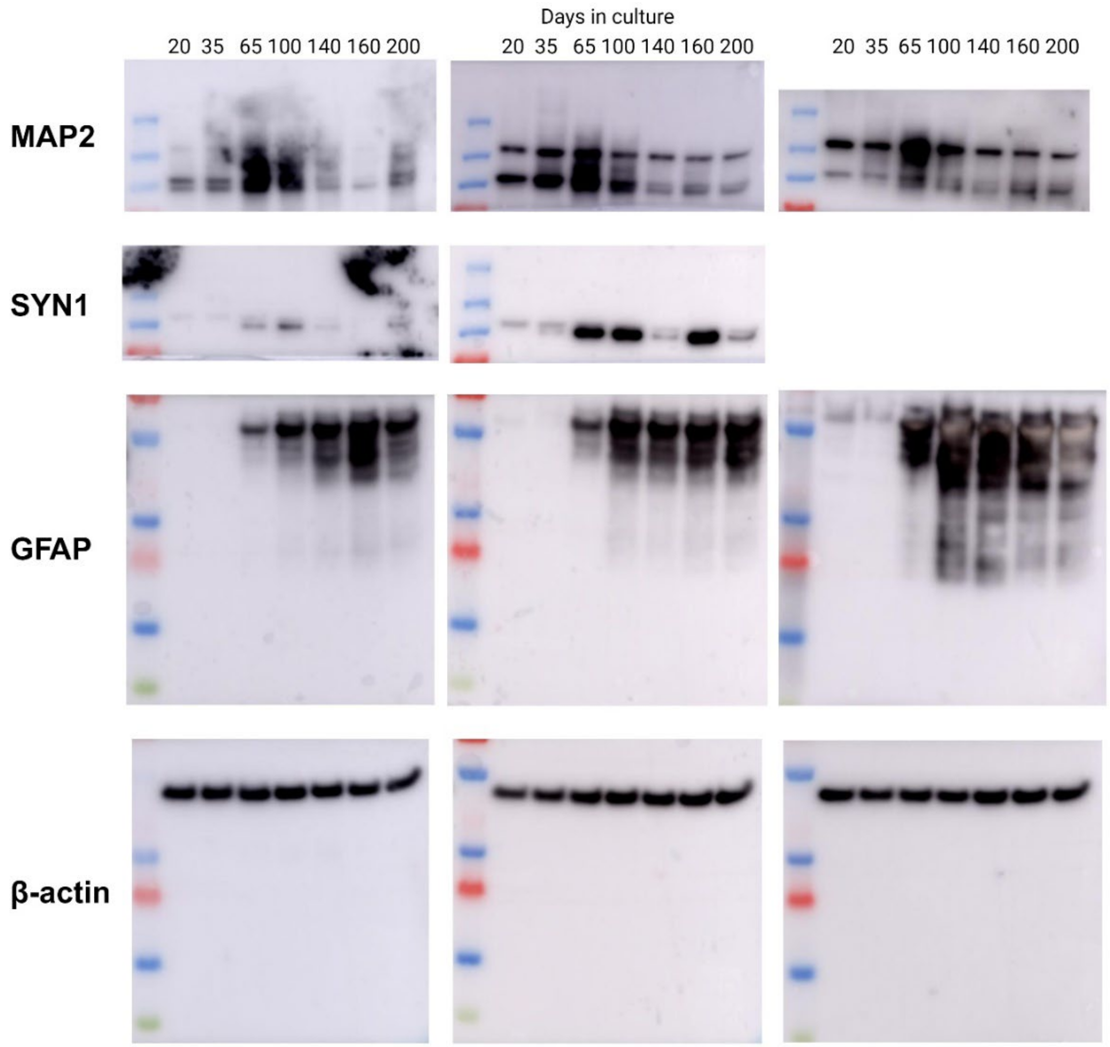


**Figure S9:** Images of Western blots of Map2a+b, synapsin 1 (SYN1), glial fibrillary acidic protein (GFAP), and β-actin in cerebral organoids at day 20, 35, 65, 100, 140, 160, and 200.

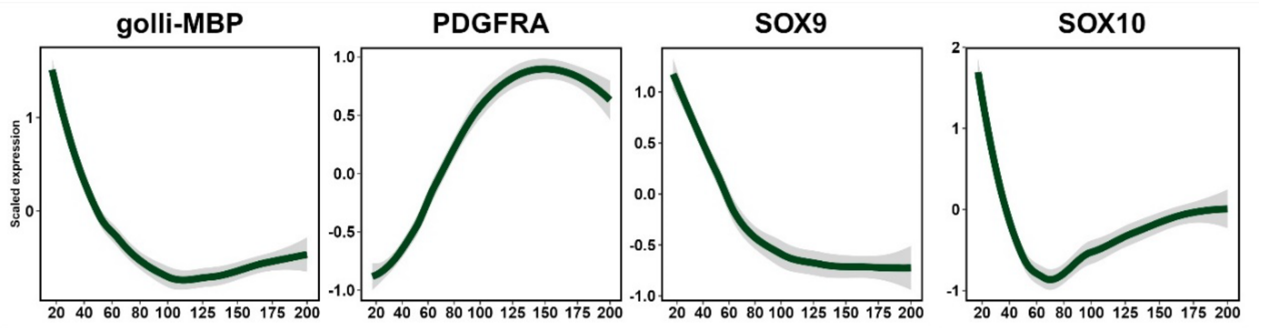


**Figure S10:** Average z-scaled abundance profiles of markers for oligodendrocyte and/or oligodendrocyte precursor cells. The average z-scaled abundances were fitted to a LOESS (locally estimated scatterplot smoothing) regression model.

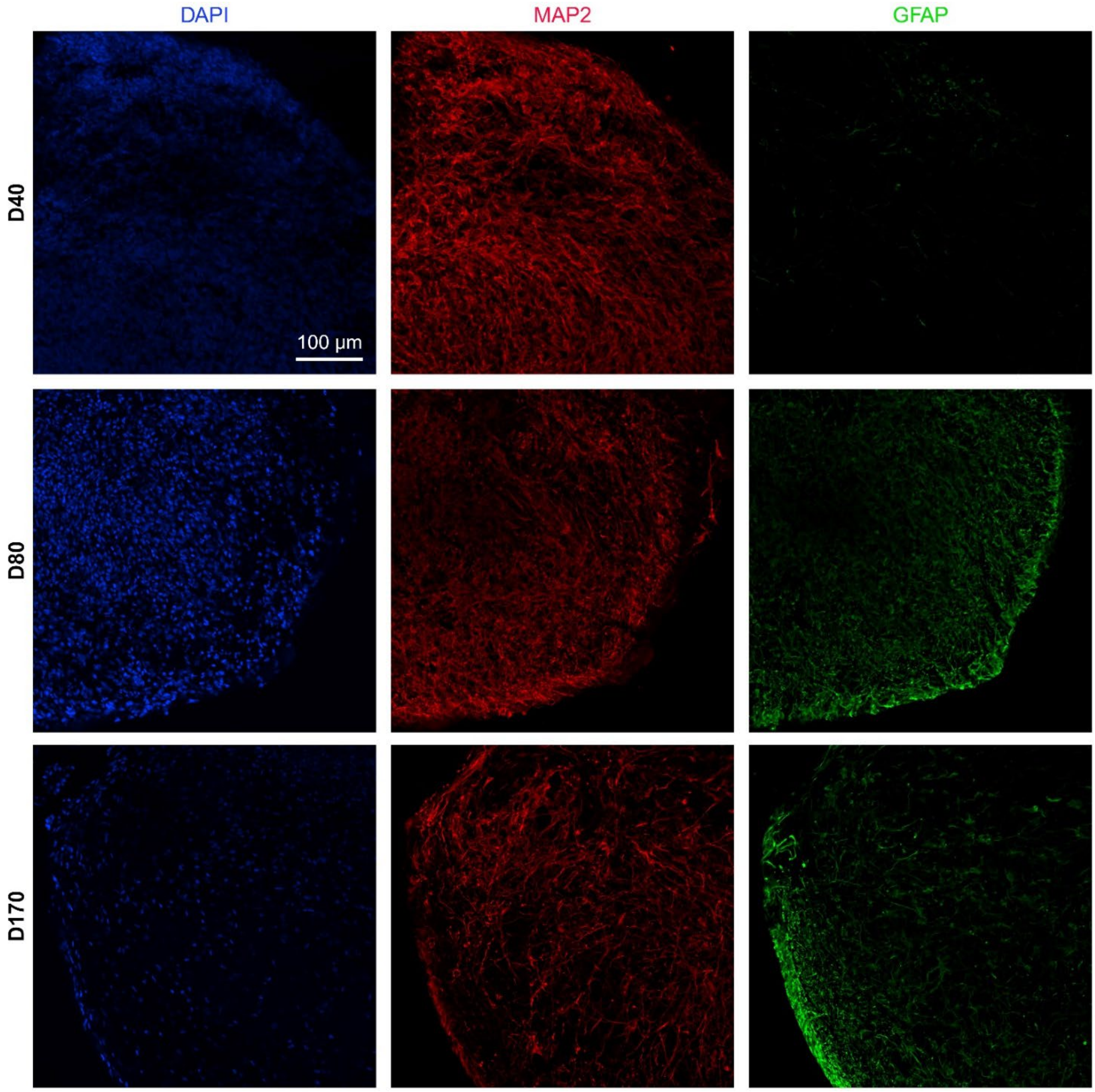


**Figure S11:** Immunohistochemistry for DAPI (blue), MAP2 (red), and GFAP (green) in cerebral organoids at day 40 (upper), 80 (middle), and 170 (lower). Scalebar: 100 µm.
